# Opposing intrinsic and synaptic plasticity mechanisms stabilize altered cortical networks during sleep deprivation

**DOI:** 10.64898/2026.05.31.729031

**Authors:** Richard J. Burman, Paul Brodersen, Hannah Alfonsa, Vladyslav V. Vyazovskiy, Colin J. Akerman

## Abstract

Sleep deprivation alters brain activity and impairs performance, yet many aspects of behaviour are preserved in this state. In the cortex for example, sleep deprivation increases neuronal firing rates and low-frequency oscillatory activity, but cortical circuits continue to process information. Recent work has implicated synaptic inhibition in these changes, with sleep deprivation causing cortical GABA_A_ receptor (GABA_A_R) signalling to become depolarizing due to changes in chloride gradients that determine the GABA_A_R reversal potential (E_GABAAR_). The impact in the intact brain remains unclear, however, as both the degree and effects of E_GABAAR_ changes depend on network activity and intrinsic properties of neurons. To address this, we perform *in vivo* gramicidin recordings from cortical pyramidal neurons in sleep-deprived mice. We reveal that synaptic E_GABAAR_ is depolarised to values sufficient to support bona fide excitatory GABA_A_R signalling, through combined network-dependent and cell-autonomous effects. In the face of this depolarizing drive, cortical neurons engage spike threshold adaptation mechanisms that limit their excitability. These opposing effects stabilise activity in simulations of sleep-deprived cortical networks and reproduce the increased firing rates and low-frequency oscillatory activity. This interplay between intrinsic and synaptic mechanisms may help maintain network stability during sleep deprivation, while reducing flexibility for subsequent adaptation.

## Introduction

Sleep deprivation impairs cognitive performance, producing lapses in attention and increased errors during demanding tasks^1^. Nevertheless, animals and humans deprived of sleep remain capable of performing many behaviours, albeit at reduced levels^2–6^. This suggests that sleep deprivation does not disable brain function but instead disrupts specific aspects of neural activity while preserving sufficient stability to support behaviour. Given its central role in higher-order functions, a key question is how the cortex is affected by a lack of sleep. Even relatively brief periods of deprivation during the normal sleep phase result in measurable changes in cortical activity. In humans, extended wakefulness increases cortical activity and low-frequency oscillatory activity in the electroencephalogram (EEG), particularly across delta (0.5-4 Hz) and theta (4-8 Hz) bands^7–13^. Similarly in mice and rats, sleep deprivation leads to increased cortical neuron action potential (i.e. spiking) activity and increased low-frequency EEG power^2,14–18^.

While the relationship between activity and sleep-wake history can vary according to neuron type and experiences during waking^19–22^, the increases in cortical spiking and low-frequency oscillatory activity during sleep deprivation must, at some level, be reflected in the synaptic and or intrinsic activity of cortical neurons. This idea aligns with the synaptic homeostasis hypothesis, which places changes in synaptic signalling at the centre of the relationship between wakefulness and cortical activity^15,20,23–25^. More specifically, an enhancement in excitatory glutamatergic synaptic signalling is consistent with the increased cortical activity observed during extended wakefulness, supported by evidence that wakefulness promotes stronger glutamatergic signalling in cortical pyramidal neurons^26–30^. A growing body of literature from humans and animals suggests that inhibitory GABAergic synaptic signalling also changes as a function of wakefulness, and in a manner consistent with increased activity levels in the sleep-deprived cortex^8,16,28,31,32^. Transcranial stimulation experiments reveal reduced intracortical synaptic inhibition in sleep-deprived humans^8^ and rodent studies demonstrate that wakefulness is associated with reduced inhibitory synaptic signalling onto cortical pyramidal neurons^28,31^. Indeed, recent work has revealed fundamental changes in the nature of postsynaptic GABAergic synaptic transmission, such that sleep deprivation is associated with a switch in the polarity of GABAergic signalling from hyperpolarizing to depolarizing^16,32,33^.

Fast cortical synaptic inhibition is mediated by chloride (Cl^-^) permeable GABA_A_ receptors (GABA_A_Rs), such that transmembrane Cl^-^ gradients determine the GABA_A_R reversal potential (E_GABAA_) and drive for Cl^-^ to either enter (hyperpolarize) or leave (depolarize) the postsynaptic neuron. Recordings from mouse cortical pyramidal neurons have shown that wakefulness results in depolarizing shifts in E_GABAAR_, due to changes in the contributions of Cl^-^ cotransporter proteins^16,32^. Such depolarizing shifts in E_GABAAR_ will not only weaken synaptic inhibition, but if sufficiently pronounced compared to a neuron’s spike threshold, could cause GABA to act as a bona fide excitatory neurotransmitter^34–38^. So far, however, estimates of E_GABAAR_ in the context of sleep deprivation have been limited to *ex-vivo* brain slices prepared from mice that experienced different amounts of prior wakefulness^16,32^. This is problematic because synaptic E_GABAAR_ and GABA_A_R signalling are known to be affected by neuronal activity levels *in vivo*, which influence the Cl^-^ loads that neurons experience^39,40^. The effects of GABA_A_R signalling *in vivo* will also depend on the state of the intrinsic properties of neurons. Membrane potential, membrane resistance, and spike threshold all affect how E_GABAAR_ and GABA_A_R signalling impact cortical neuronal activity^34,40^. Indeed, these intrinsic properties are themselves dynamic, providing neurons with adaptive mechanisms for either amplifying or counteracting changes in synaptic signalling^41–44^.

Here we use *in vivo* gramicidin perforated patch-clamp recordings to investigate inhibitory synaptic transmission and intrinsic properties in layer 2/3 (L2/3) cortical pyramidal neurons in sleep-deprived mice. Under these recording conditions that preserve native ion gradients, we reveal that sleep deprivation causes synaptic E_GABAAR_ to depolarize to levels that are sufficient to support excitatory GABA_A_R signalling. In the face of increased depolarizing drive however, we find that spike threshold adaptation mechanisms act to limit the excitability of cortical pyramidal neurons in the sleep-deprived state. This combination of opposing intrinsic and synaptic plasticity mechanisms is shown to stabilize overall activity in simulations of sleep-deprived cortical networks, whilst reproducing experimentally observed increases in spiking activity and low-frequency oscillatory activity.

## Results

### Synaptic E_GABAAR_ is depolarized in L2/3 pyramidal neurons during sleep deprivation

We investigated the synaptic and intrinsic properties of cortical neurons in mice that were well entrained to the light-dark cycle, spending most of the time asleep during the light (inactive) period and most of the time awake during the dark (active) period (see Methods). Recordings were performed in awake mice at Zeitgeber Time 3 (ZT3), and the mice were randomly assigned to one of two conditions that differed in terms of their recent sleep-wake history (**Fig. 1a**; see Methods). Mice in the rested condition were allowed to follow their normal sleep-wake pattern, such that the 3-h period prior to recording (ZT0-ZT3) was dominated by sleep (see Methods)^16^. Mice in the sleep-deprived (SD) condition meanwhile, were subjected to an established sleep deprivation (SD) protocol during the 3-h period between ZT0-ZT3, which consisted of exposing the animal to novel objects (**Fig. 1a**; see Methods). This SD protocol has been shown to result in increased sleep pressure that is evident in both the awake state and during subsequent sleep^16,32^.

**Figure 1:**
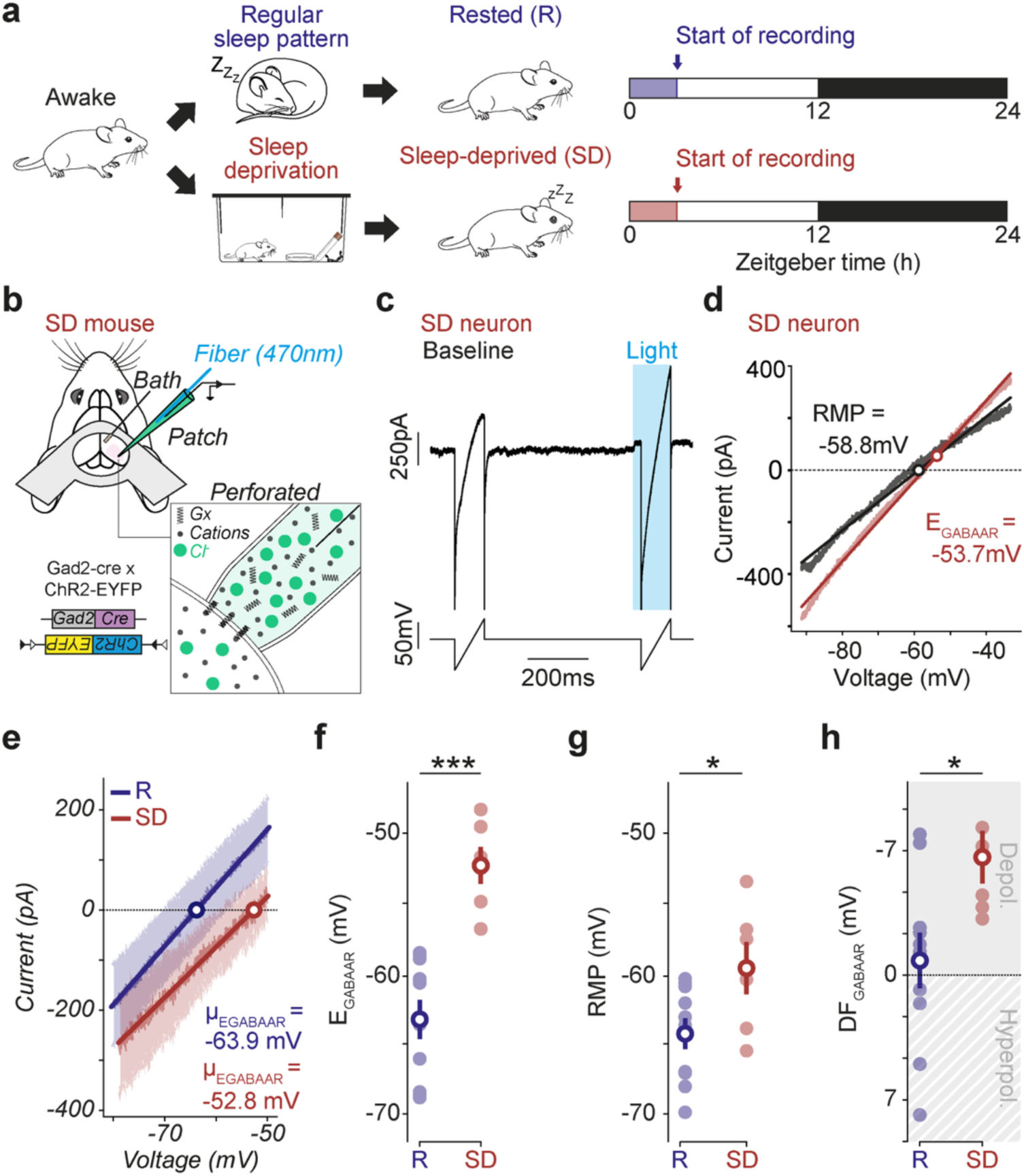
Synaptic E_GABAAR_ is depolarized in L2/3 cortical pyramidal neurons during sleep deprivation. **a**, Mice expressing ChR2 in Gad2-positive interneurons were either rested (R, blue) as part of their regular sleep pattern or underwent an acute sleep deprivation protocol between ZT0 and ZT3 (SD, red). Recordings were performed at ZT3-ZT6, while animals were awake. **b**, *In vivo* gramicidin perforated patch-clamp recordings were performed from L2/3 pyramidal neurons in V1. Presynaptic GABA release was initiated from nearby interneurons by delivering blue light (470 nm) pulses via the patch pipette. **c**, Voltage-clamp recording from a neuron in a SD mouse, consisting of a voltage ramp under control conditions (Baseline) and a voltage ramp during the light-evoked postsynaptic GABA_A_R conductance (Light). **d**, Current-voltage plot of baseline (black) and light ramp current (red) from the neuron in ‘c’. RMP, resting membrane potential. **e**, Population plot of synaptic GABA_A_R currents recorded in the rested (n = 10 neurons, 9 mice) and SD conditions (n = 6 neurons, 6 mice). Line indicates mean, shading indicates SEM. **f**, E_GABAAR_ was more depolarized in the SD compared to rested condition (R: -63.3 ± 1.4 mV vs. SD: -52.3 ± 1.3 mV, p < 0.001, *t-test*). **g**, RMP was more depolarized in the SD compared to resting condition (R: -64.2 ± 1.1 mV vs. SD: -59.4 ± 1.9 mV, p = 0.04, *t-test*). **h**, DF_GABAAR_ was more depolarized in the SD compared to resting condition (R: -0.9 ± 1.7 mV vs. SD: –7.1 ± 1.6mV, p = 0.03, *t-test*). DF_GABAAR_ in the rested condition was not different to zero, consistent with an overall shunting effect (p = 0.61, *one-sample t-test*). In contrast, DF_GABAAR_ was depolarizing in the SD condition (p = 0.006, *one-sample t-test*). ns, non-significant; p > 0.05; *, p < 0.05; ***, p < 0.001.

To determine the extent to which synaptic GABA_A_R signalling is altered in SD cortex, we performed *in vivo* gramicidin perforated patch clamp recordings from L2/3 pyramidal neurons in primary visual cortex (V1) of head-fixed mice (**Fig. 1b** and **Supplementary Fig. 1**). The gramicidin recording method leaves transmembrane Cl^-^ gradients intact, making it possible to determine the native reversal potential of GABA_A_Rs (E_GABAAR_)^39^. To selectively activate GABAergic synaptic inputs, we used mice expressing channelrhodopsin-2 (ChR2) in Gad2-positive interneurons. Brief pulses of blue light were delivered through an optic fiber within the recording pipette, eliciting GABA release from local presynaptic terminals (**Fig. 1b**). Meanwhile, postsynaptic GABA_A_R-responses in the pyramidal neurons were recorded in voltage-clamp mode using a ramp protocol. This allowed us to generate current-voltage (IV) plots, from which the resting membrane potential (RMP) and E_GABAAR_ could be determined (**Fig. 1c–d**).

Compared to rested mice, synaptic E_GABAAR_ was found to be significantly more depolarized in L2/3 pyramidal neurons in SD mice (**Fig. 1e-f**), with a mean value (-52.3 ± 1.3 mV) that is close to reported values for the spike threshold in cortical pyramidal neurons, suggesting that GABA_A_Rs could exert excitatory effects in the SD state^41,45,46^. Indeed, the shift in **E_GABAAR_** was accompanied by a more modest shift in RMP (**Fig. 1g**), such that when the driving force for synaptic GABA_A_R currents was computed (DF_GABAAR_; defined as RMP - E_GABAAR_), we observed a consistent depolarizing shift in the SD condition (**Fig. 1h**). Synaptic DF_GABAAR_ in rested animals was close to zero, indicating that GABAergic inputs are neither strongly hyperpolarizing nor depolarizing in this state, in line with a shunting role for GABA_A_R signalling in the rested cortex^39,40^. In contrast, synaptic DF_GABAAR_ in SD animals was consistently depolarizing, indicating that GABA could operate as an excitatory transmitter in the SD cortex.

### Network activity contributes to depolarized synaptic E_GABAAR_ during sleep deprivation

Previous theoretical and experimental work suggests that the current level of network activity can itself affect **E_GABAAR_**. For example, more active network states are associated with depolarized membrane potentials (V_m_) and greater activation of GABA_A_R-containing synapses, which increase the Cl^-^ loads experienced by postsynaptic neurons^39,40,47^. Consistent with reports of increased network activity in the SD cortex^15,48,8,7^, our current-clamp recordings revealed that L2/3 pyramidal neurons in SD mice exhibited more depolarized mean V_m_ values and greater fluctuations in subthreshold V_m_, which reflect levels of synaptic activity (**Supplementary Fig. 2**)^39^. To determine the contribution of current network activity to the depolarized **E_GABAAR_** in SD cortex, we silenced local synaptic activity by injecting the AMPA receptor antagonist NBQX during our *in vivo* gramicidin recordings (**Fig. 2a**). NBQX administration led to a pronounced hyperpolarization of V_m_ and almost complete suppression of subthreshold V_m_ fluctuations (**Fig. 2b** and **Supplementary Fig. 3**), confirming the effective silencing of local network activity. This allowed for within-neuron comparisons to test the prediction that the levels of network activity contribute to the depolarized synaptic E_GABAAR_ observed in SD (**Fig. 2c**).

**Figure 2:**
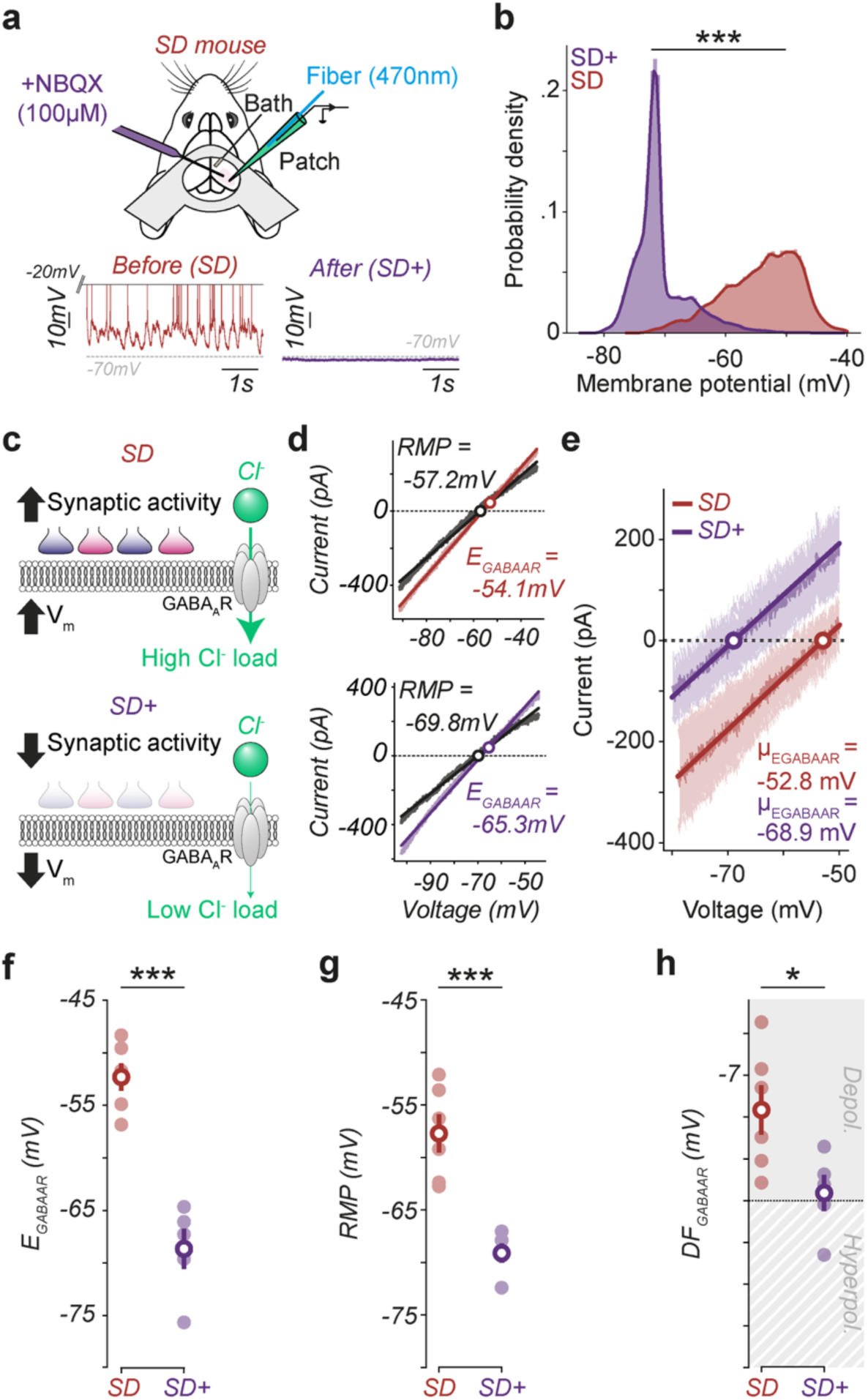
Network activity during sleep deprivation contributes to the depolarized synaptic E_GABAAR_. **a**, Gramicidin perforated patch-clamp recordings were performed from L2/3 pyramidal neurons in SD mice and network activity was silenced by local NBQX delivery. Example current-clamp recordings before (SD, red) and after NBQX (SD+, purple). **b**, V_m_ values differed before (n = 9 cells, 9 mice) and after NBQX (n = 5 cells, 5 mice; p < 0.001, *Kolmogorov–Smirnov test*). **c**, Illustration showing how depolarized V_m_ and synaptic activity are conducive to GABA_A_R-mediated Cl^-^ influxes (top), and how this is reduced upon silencing network activity (bottom). **d**, Current-voltage plots from a neuron before (SD, top) and after NBQX (SD+, bottom). **e**, Population plot of synaptic GABA_A_R currents recorded before (SD, n = 6 cells, 6 mice) and after NBQX (SD+, n = 5 cells, 5 mice). **f**, Silencing network activity caused a hyperpolarizing shift in E_GABAAR_ (SD: -52.3 ± 1.3 mV vs. SD+: -68.6 ± 1.9 mV, p < 0.001, *t-test*). **g**, Silencing network activity caused a hyperpolarizing shift in RMP (SD: -57.8 ± 1.8 mV vs. SD+: -69.1± 0.9 mV, p = 0.0005, *t-test*). **h**, Silencing network activity caused a hyperpolarizing shift in synaptic DF_GABAAR_ (SD: -5.4 ± 1.5 mV vs. SD+: -0.5 ± 0.9 mV, p = 0.02, *t-test*), from depolarizing (p = 0.01, *one-sample t-test*) to shunting (p = 0.7, *one-sample t-test*). *, p <0.05, ***, p < 0.001.

Voltage ramp protocols before and after NBQX delivery revealed a pronounced hyperpolarizing shift in synaptic E_GABAAR_ within the same neuron (**Fig. 2d**). Population analysis confirmed this, showing a consistent and robust hyperpolarization of synaptic E_GABAAR_ following the silencing of network activity (**Fig. 2e-f**). These changes were accompanied by a less pronounced hyperpolarizing shift in RMP (**Fig. 2g**), resulting in a net hyperpolarizing shift in DF_GABAAR_ (**Fig. 2h**). Together these data support the conclusion that the current network activity in the SD state contributes to the depolarized synaptic E_GABAAR_ observed in SD cortex, likely through ongoing Cl^-^ influxes via GABA_A_Rs under conditions of high synaptic activity and depolarized V_m_^39,40^.

### Cell-autonomous changes underpin depolarized synaptic E_GABAAR_ during sleep deprivation

*In vitro* recordings have indicated that a period of SD can lead to a depolarizing shift in E_GABAAR_ due to changes in a neuron’s Cl^-^ cotransporter proteins^16,33,39^. To establish whether such cell-autonomous effects underlie the depolarized E_GABAAR_ observed in SD cortex, we compared E_GABAAR_ in rested and SD mice having eliminated local network activity in both conditions (**Fig. 3a**). We first confirmed that silencing network activity with NBQX was equally effective in rested and SD mice, such that V_m_ was hyperpolarized to comparable levels and subthreshold V_m_ fluctuations showed almost complete suppression (**Supplementary Fig. 4**). Under these matched recording conditions, voltage ramp protocols continued to reveal a clear difference in E_GABAAR_ between neurons in rested and SD mice (**Fig. 3b-c**). E_GABAAR_ remained more depolarized in neurons from SD mice compared to rested mice (**Fig. 3d**), RMP was not different (**Fig. 3e**), and DF_GABAAR_ remained more depolarized in the SD condition (**Fig. 3f**). These data support the conclusion that cell-autonomous mechanisms underlie the differences in E_GABAAR_ between rested and SD. Therefore, the depolarized E_GABAAR_ observed in SD cortex *in vivo* is a consequence of both the state of network activity and underlying cell-autonomous changes in Cl^-^ regulatory mechanisms^16,39^.

**Figure 3:**
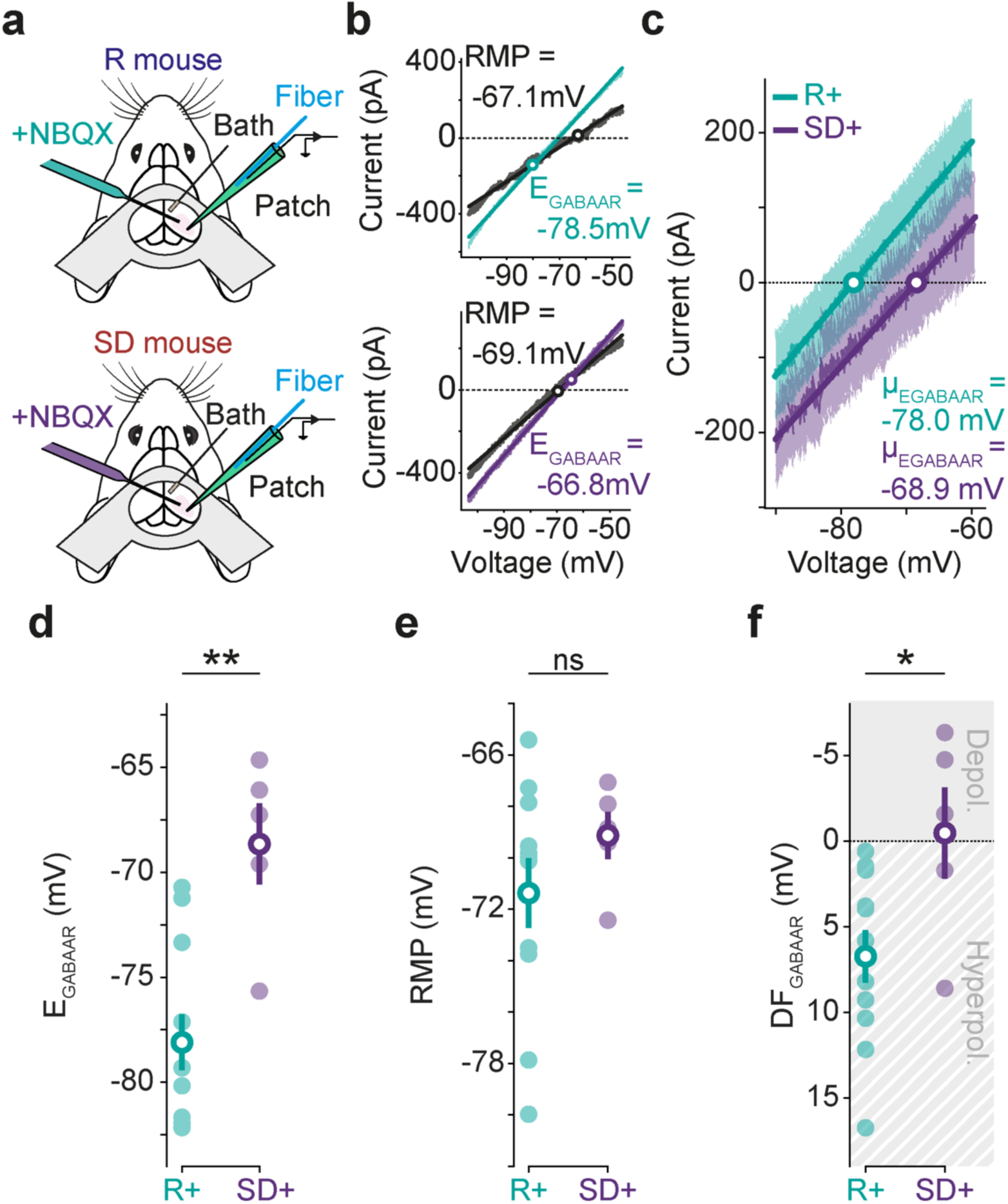
Cell-autonomous changes underpin the depolarized synaptic E_GABAAR_ in sleep-deprived cortex. **a**, Network activity was silenced by local delivery of the glutamate receptor blocker, NBQX, and gramicidin perforated patch-clamp recordings were performed from L2/3 pyramidal neurons in rested (R, blue) or sleep-deprived (SD, red) mice. **b**, Current-voltage plots from voltage ramp protocols performed on a neuron in a network-silenced rested mouse (R+, turquoise, top), and in a network-silenced SD mouse (SD+, purple, bottom). **c**, Population plot of synaptic GABA currents recorded in network-silenced rested mice (R+, n = 11 neurons, 7 mice) and network-silenced SD mice (SD+, n = 5 neurons, 5 mice). **d**, Network-silenced E_GABAAR_ was more depolarized in the SD condition (R+: -78.1 ± 1.3 mV vs. SD+: -68.7 ± 1.9 mV, p = 0.001, *t-test*). **e**, Network-silenced RMP was not different between SD and rested conditions (R+: -71.4 ± 1.4 mV vs. SD+: -69.1± 0.9 mV, p = 0.3, *t-test*). **e**, Network-silenced DF_GABAAR_ was more depolarized in the SD condition (R+: 6.7 ± 1.5 mV vs. SD+: -0.5 ± 2.7 mV, p = 0.03, *t-test*). ‘ns’, non-significant (p > 0.05); *, p < 0.05; **, p < 0.01.

### Intrinsic neuronal plasticity mechanisms oppose increased depolarizing drive during sleep deprivation

Our results establish that L2/3 pyramidal neurons in SD cortex are characterized by their depolarized state, both in terms of their GABA_A_R-mediated synaptic inputs and subthreshold V_m_. Such pronounced depolarizing effects would be expected to result in hyperexcitability, yet neurons in SD cortex are not associated with uncontrolled spiking activity^15,16^. We therefore hypothesised that intrinsic neuronal plasticity mechanisms act to compensate for the increased depolarising drive during SD. To test this possibility, we performed gramicidin current-clamp recordings of periods of spontaneous activity from L2/3 pyramidal neurons and then analysed the subthreshold V_m_ and action potential properties (**Fig. 4a-b**). These data revealed that the range and distribution of subthreshold V_m_ values was more depolarized in SD mice than in rested mice (**Fig. 4c**), consistent with our observation that mean E_GABAAR_ and mean V_m_ are both more depolarized (**Fig. 1** and **Supplementary Fig. 2**). To compare action potential activity between the rested and SD cortex, we identified individual action potentials during the spontaneous recordings, generated phase plots based on the first derivative of V_m_ and used these to define action potential thresholds (**Fig. 4d**). This revealed that the threshold for spike generation was significantly more depolarized in neurons from SD cortex (**Fig. 4e**), consistent with the engagement of spike threshold adaptation mechanisms^41,42,49–51^. If neurons in SD cortex engage spike threshold adaptation mechanisms in response to the increased depolarizing drive, we would predict that spike threshold would reflect both a neuron’s absolute V_m_ value and the rate of V_m_ change before action potential initiation^41,42,49^. In line with these predictions, we observed that spike threshold was strongly positively correlated with the absolute V_m_ value immediately prior to spike initiation, and strongly negatively correlated with the rate of membrane depolarization immediately prior to spike initiation (**Fig. 4f**). These data support the conclusion that under the conditions of increasing depolarizing drive, neurons in SD cortex engage intrinsic plasticity mechanisms that raise the effective spike threshold.

**Figure 4:**
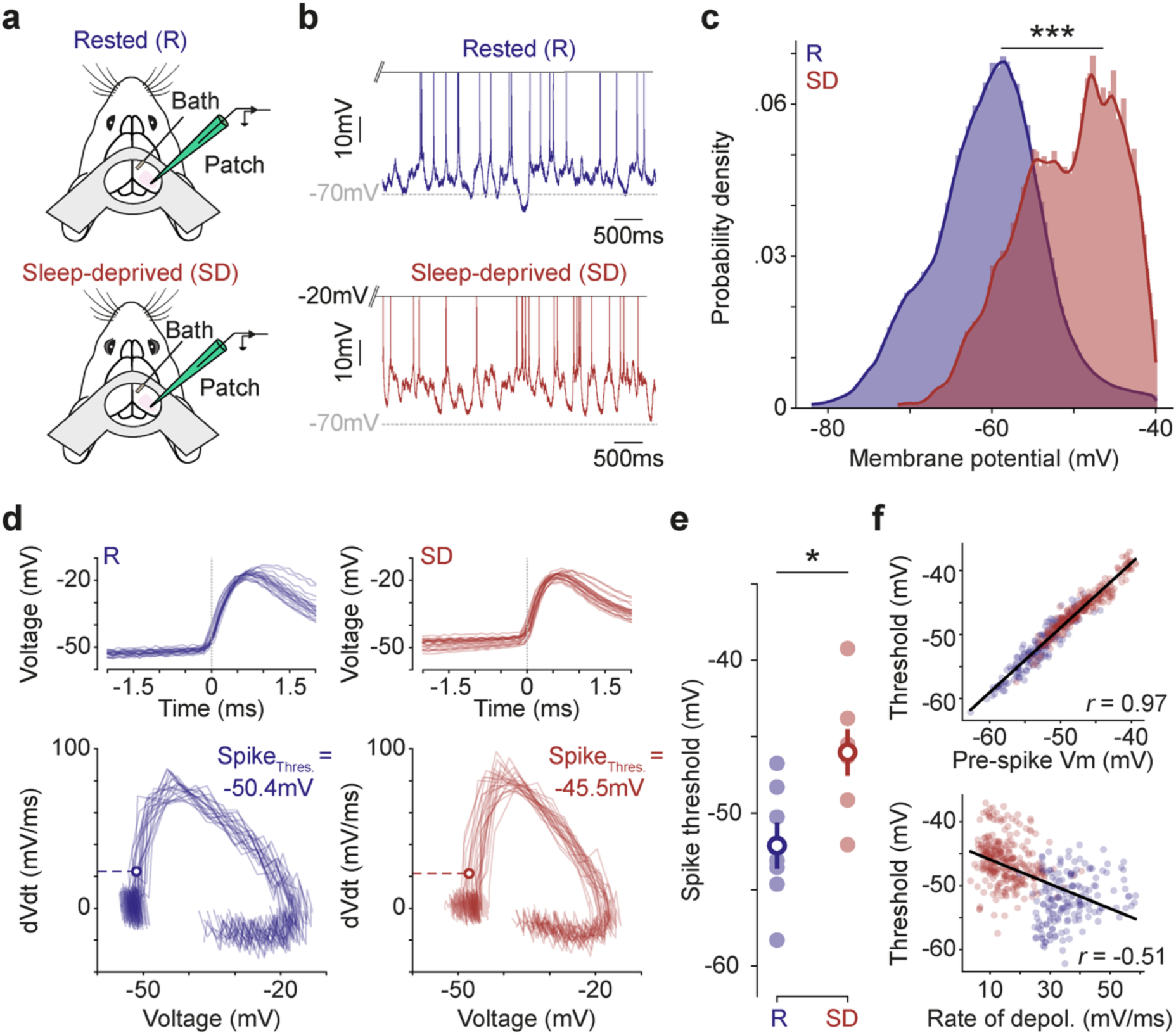
Sustained membrane depolarization engages spike threshold adaptation mechanisms in the sleep-deprived cortex. **a**, Current-clamp recordings were performed to quantify membrane potential (V_m_) dynamics in L2/3 pyramidal neurons in rested (R, blue) or sleep-deprived (SD, red) mice. **b**, Example recordings show subthreshold V_m_ and spiking activity of a neuron from a rested (top) and a SD mouse (bottom). **c**, V_m_ values differed between rested (n = 12 neurons, 10 mice) and SD (n = 9 neurons, 9 mice; p < 0.001, *Kolmogorov–Smirnov test*) conditions. **d**, Action potential plot (top) and corresponding phase plot (bottom) for a neuron from a rested (left) and SD (right) mouse. Spikes recorded over a period of 5 s are overlaid. Phase plots show rate of change in V_m_ (dV/dt) as a function of V_m_. Spike threshold (Spike_Thres_) is indicated with an empty circle on the phase plots and a grey vertical line on the action potential plots. **e**, Spike threshold was more depolarized in neurons from the SD condition (R: -52.1 ± 1.5 MΩ n = 7 neurons, 7 mice vs. SD: -46.1 ± 1.5 MΩ, n = 7 neurons, 7 mice, p = 0.02, *t-test*). *, p < 0.05, ***, p < 0.001. **f**, Spike threshold exhibited a strong positive correlation with the mean V_m_ immediately before spike (top; r = 0.97, p < 0.001) and a strong negative correlation with the rate of V_m_ depolarization immediately before spike (bottom; r = -0.51, p < 0.001).

### Sleep-deprived cortex exhibits increased in pyramidal neuron spiking and low-frequency oscillatory activity

To investigate how the opposing changes in intrinsic neuronal and synaptic properties interact with one another in SD cortex, we quantified *in vivo* network activity in two ways. First, mean firing rates of L2/3 pyramidal neurons were analysed from the periods of spontaneous activity recorded during the gramicidin perforated-patch current-clamp recordings in head-fixed mice (**Fig. 5a**). These recordings revealed regular spiking activity in both the rested and SD condition, with neurons in SD mice displaying approximately two-fold higher mean firing rates (**Fig. 5b-c**), consistent with previous reports^15^. Second, to assess slower network dynamics over longer temporal windows, we analysed continuous cortical local field potential (LFP) recordings from freely-moving mice (**Fig. 5d**). This enabled us to compare a rested LFP during a mouse’s normal sleep- wake pattern, with a SD LFP recorded when the mouse was subjected to the same SD protocol used in the head-fixed recordings (**Fig. 5d**; see Methods). Power spectral analysis of the LFP revealed a clear increase in power at frequencies below 20 Hz in the SD condition compared to the rested condition (**Fig. 5e**), which was most pronounced in the delta frequency band (1-4 Hz; **Fig. 5e** and **Supplementary Fig. 5**). Such an increase in low-frequency oscillatory activity is characteristic of drowsy or “sleep-like” cortical states, and has previously been reported in animals and humans following SD^2,3,16,52^. Therefore, the SD cortical network is more active overall, and its activity is consistent with increased synchronisation, particularly at lower frequencies.

**Figure 5:**
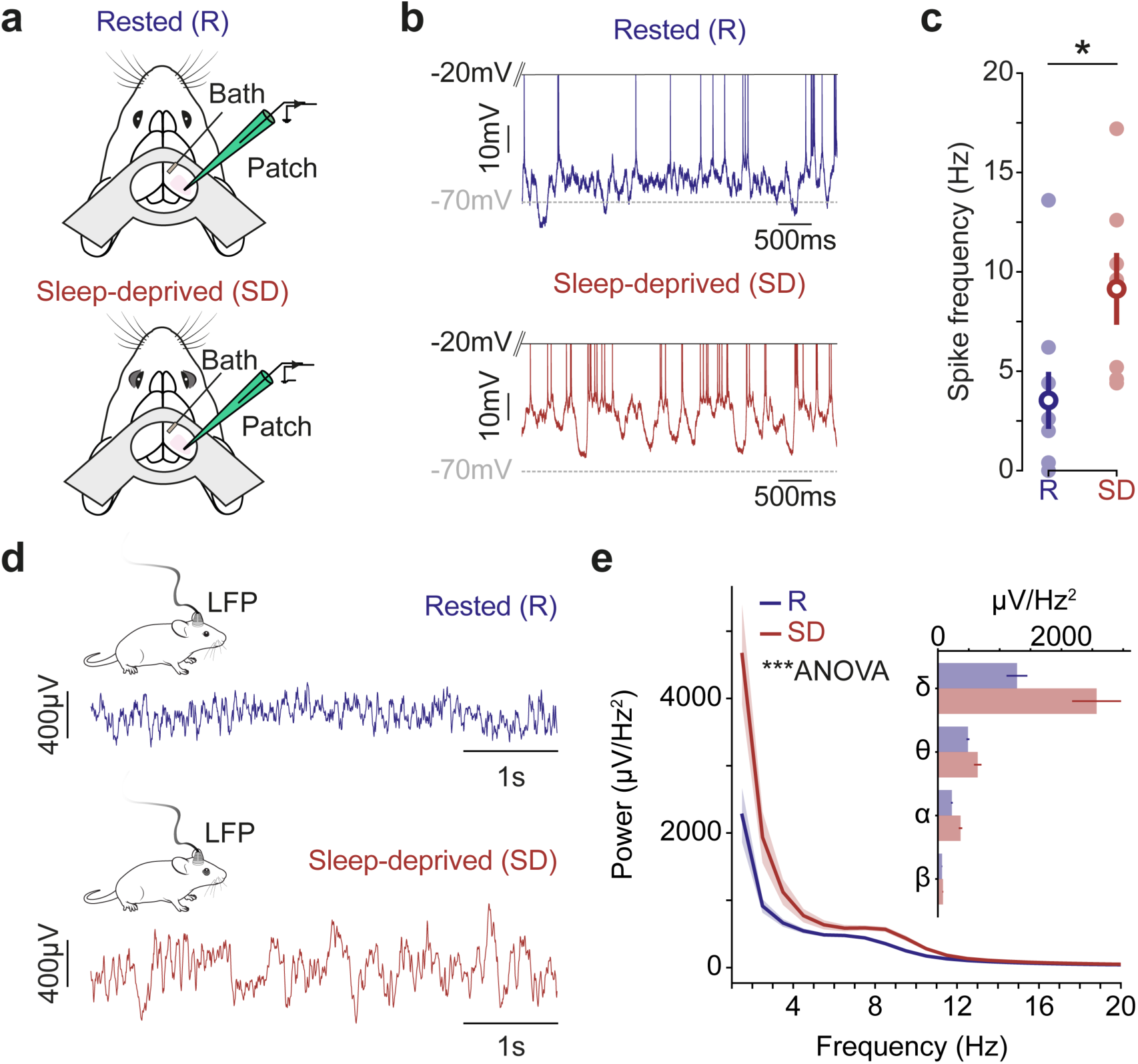
Sleep-deprived cortex exhibits increased pyramidal neuron spiking and low-frequency oscillatory activity in the local field potential. **a**, Current-clamp recordings were used to quantify spontaneous spiking activity in L2/3 pyramidal neurons in rested (R, blue) or sleep-deprived (SD, red) mice. **b**, Example recordings show spiking activity in a rested (top) and SD (bottom) mouse. **c**, Spike frequency was higher in neurons from the SD condition (R: 3.5 ± 1.4 Hz n = 9 neurons, 9 mice vs. SD: 9.1 ± 1.8 Hz, n = 7 neurons, 7 mice, p = 0.03, *unpaired t-test*). **d**, Example local field potential (LFP) recordings from the same freely-moving awake mouse, which had either rested (top) or experienced SD during the preceding ZT0-ZT3 (bottom). **e**, LFP power spectra reveal increased low-frequency oscillatory activity in the SD compared to rested condition (n = 22 mice, p < 0.001, *two-way repeated measures Anova*). Line indicates mean, shading indicates SEM. Inset plot shows mean power in the delta (1-4 Hz), theta (4-8 Hz), alpha (8-12 Hz), and beta (12-20 Hz) frequency bands. *, p < 0.05; ***, p < 0.001.

### Opposing effects of spike threshold adaptation and depolarized E_GABAAR_ can generate the changes in cortical activity observed during sleep deprivation

To explore whether the opposing effects of spike threshold adaptation and depolarized E_GABAAR_ could account for the observed changes in cortical network activity, we constructed a biophysically-informed spiking network model composed of excitatory pyramidal neurons and inhibitory interneurons (**Fig. 6a**; see Methods). Model parameters for resting membrane potential, E_GABAAR_, and spike threshold were taken directly from our experimental measurements under rested and SD conditions, and each neuron received a mixture of independent and shared excitatory input. To describe activity in the model we measured the spiking output of the pyramidal neurons and used the power spectrum of excitatory synaptic inputs as an estimate of the LFP^53^. The pyramidal neurons exhibited regular spiking activity in simulations of both the rested and SD cortex, but mean firing rates were consistently higher in the simulated SD condition (**Fig. 6b-c**). Meanwhile, estimates of the LFP revealed an increase in power at frequencies below 20 Hz in the SD condition, which was greatest in the delta frequency band (1-4 Hz; **Fig. 6d-e** and **Supplementary Fig. 6**). These findings therefore recapitulated key features of our *in vivo* observations, indicating that elevated spiking and enhanced low-frequency oscillatory activity in the SD cortex can be accounted for by the opposing changes in E_GABAAR_ and spike threshold.

**Figure 6:**
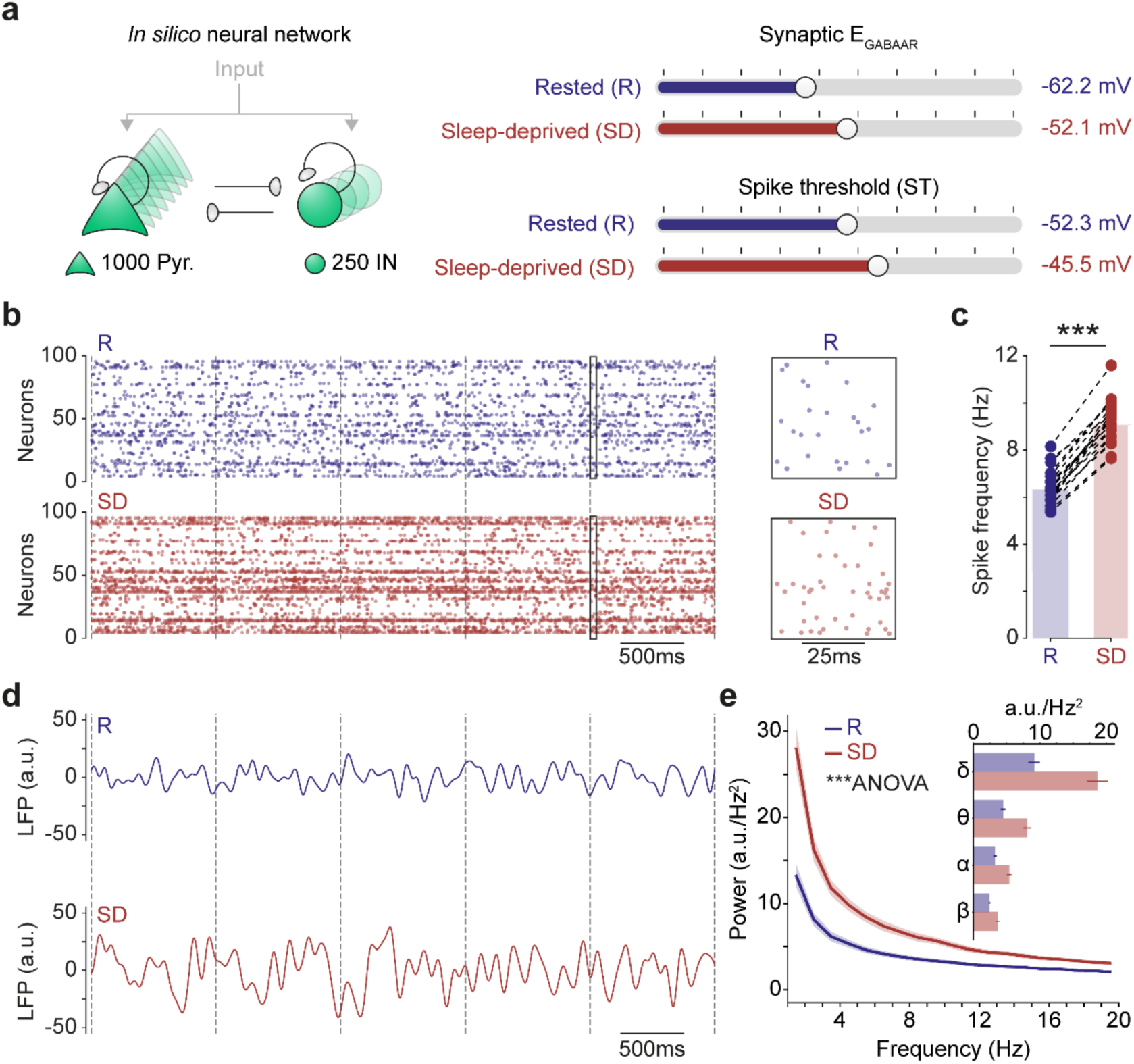
Spike threshold adaptation and depolarized E_GABAAR_ increase spiking and low-frequency oscillatory activity in a simulated sleep-deprived network. **a**, Schematic of network model (left) consisting of interconnected excitatory pyramidal neurons (Pyr.) and inhibitory interneurons (IN). Synaptic E_GABAAR_ and spike threshold in the pyramidal neurons were set to the mean values observed experimentally (right) under rested (R, blue) and sleep-deprived (SD, red) conditions. **b**, Raster plots showing the spiking activity of 100 pyramidal neurons in a simulated rested (top) and SD network (bottom). Insets show a 50 ms period from the raster plots. **c**, Pyramidal neuron spiking was higher in the simulated SD network (R: 6.4 ± 0.2 Hz vs. SD: 9.2 ± 0.2 Hz, n = 20 simulations in each condition, p < 0.001, *Wilcoxon paired test*). **d**, Example LFPs from a simulated rested (top) and SD network (bottom). **e**, LFP power spectra reveal increased low-frequency oscillatory activity in the SD network compared to the rested network (n = 20 simulations per condition, p < 0.001, *two-way repeated measures Anova*). Line indicates mean, shading indicates SEM. Inset plot shows mean power in different frequency bands. ***, p < 0.001.

### Opposing effects of spike threshold adaptation and depolarized E_GABAAR_ stabilise activity in a sleep-deprived network

To dissect the contributions of spike threshold adaptation and depolarized E_GABAAR_, we simulated two additional, hypothetical SD network conditions (**Fig. 7a**). In the first of these, we prevented intrinsic neuronal plasticity so that the spike threshold maintained the same value as observed in rested mice. In the second hypothetical SD network, we removed the depolarizing effects of E_GABAAR_, by fixing this at the more hyperpolarized value observed in rested mice. Strikingly, the simulated SD network without spike threshold adaptation exhibited dramatically elevated pyramidal neuron spiking activity, nearly 20-fold higher than the control SD condition (**Fig. 7b-c**), and reminiscent of firing rates observed during epileptiform discharges^54,55^. Meanwhile, the simulated SD network without the E_GABAAR_ shift exhibited a marked reduction in pyramidal neuron firing, falling to one-third of that observed in control SD simulations (**Fig. 7b-c**). Power analysis of the simulated LFP revealed that both hypothetical networks exhibited an almost complete collapse in oscillatory power below 20 Hz (**Fig. 7d-e**)^2,16–18^. The SD network without spike threshold adaptation was dominated by very high-frequency oscillatory activity (∼ 200 Hz), as a result of the dramatic increase in spiking activity, and again consistent with recordings during epileptiform discharges^56^ (**Fig. 7e**). Meanwhile, due to its lower spiking activity, the SD network without E_GABAAR_ shift exhibited a substantial reduction in power below 20 Hz frequency range, also contrary to what is observed experimentally (**Fig. 7e**). These findings support the conclusion that the opposing synaptic and intrinsic neuronal changes are both required to generate the SD network phenotype. Depolarized E_GABAAR_ promotes the recruitment of cortical neurons to increase synchronous spiking activity, while spike threshold adaptation prevents runaway excitation and maintains stable network activity.

**Figure 7:**
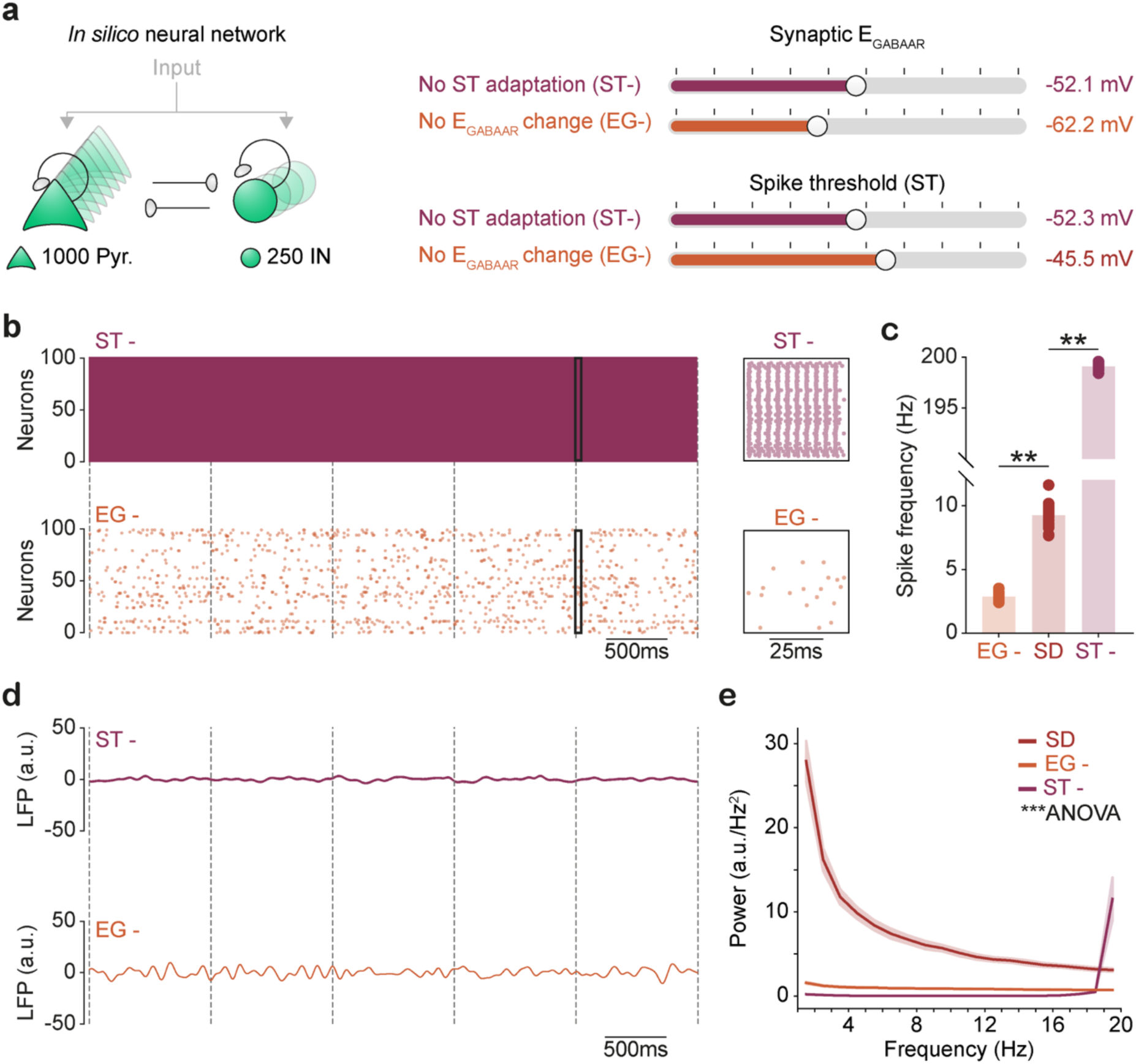
Opposing effects of spike threshold adaptation and depolarized E_GABAAR_ stabilise activity in a simulated sleep-deprived network. **a**, Network model (left) in which pyramidal neurons were assigned different combinations of experimentally observed values of synaptic E_GABAAR_ and spike threshold (ST) to simulate hypothetical states of a sleep-deprived (SD) network. The “No ST adaptation” (ST-, burgundy) network used the E_GABAAR_ value from SD animals and ST adaptation from R animals. The “No E_GABAAR_ change” (EG-, orange) network used the ST value from SD animals and the E_GABAAR_ from R animals. **b**, Raster plots showing spiking of 100 pyramidal neurons in ST- network (top) or EG- network (bottom). Insets show a 50 ms period from the raster plots. **c**, Compared to the control SD network, pyramidal neuron spiking was 20-fold higher without ST adaptation (SD: 9.2 ± 0.2 Hz vs. ST-: 199.1 ± 0.1 Hz, n = 20 in each condition, p = 0.005, *Dunn’s test*) and 3-fold lower without the depolarizing shift in E_GABAAR_ (SD: 9.2 ± 0.2 Hz vs. EG-: 2.9 ± 0.1 Hz, n = 20 in each condition, p = 0.005, *Dunn’s test*). **d**, Example LFPs from a ST- network (top) or EG-network (bottom). **e**, Compared to a control SD network (SD, red), there was a 10-fold power-decrease in low-frequency oscillatory activity in the SD network without ST adaptation (ST-, burgundy; n = 20 simulations per condition, p < 0.001, *two-way repeated measures Anova*) and the SD network without E_GABAAR_ change (EG-, orange; p < 0.001, *two-way repeated measures Anova*). **, p < 0.01; ***, p < 0.001.

## Discussion

To investigate GABA_A_R signalling and intrinsic excitability in the sleep-deprived cortex, we studied layer 2/3 pyramidal neurons using *in vivo* gramicidin perforated patch-clamp recordings. Under conditions that preserve ionic driving forces, we reveal that synaptic E_GABAAR_ becomes markedly depolarized in sleep-deprived cortex, reflecting contributions from both the current network activity and underlying cell-autonomous mechanisms. The net effect would predict excitatory actions of GABA_A_R signalling in the sleep-deprived cortex. However, we reveal that in the face of the depolarizing drive, pyramidal neurons engage spike threshold adaptation mechanisms that oppose the depolarizing effects of synaptic E_GABAAR_, and limit spiking activity. In biologically-relevant simulations of the sleep-deprived cortex, the combined effect of these opposing mechanisms is a stable network state that is characterised by increased pyramidal neuron spiking activity and an increase in low-frequency oscillatory activity, both of which are observed experimentally.

Our findings support a central role for altered GABA_A_R signalling, consistent with previous work that has detected decreases in intracortical inhibition following a period of sleep deprivation in animals and humans^16,32,57,58^. In providing the first synaptic E_GABAAR_ measurements in the intact sleep-deprived cortex, we reveal that E_GABAAR_ reaches values that are close to typical spike thresholds in cortical pyramidal neurons^41,45^. This suggests that GABA could operate as an excitatory neurotransmitter, particularly given how temporally and spatially isolated synaptic inputs are integrated by pyramidal neurons^46^. The depolarized E_GABAAR_ values in sleep-deprived cortex are shown to reflect both the ongoing *in vivo* network activity and cell-autonomous changes that are independent of network activity. The contribution of network activity was comparable in magnitude between the rested and sleep-deprived cortex, suggesting that neurons in both conditions experience similar Cl^-^ loads, potentially reflecting limits on the extracellular GABA levels responsible for phasic and tonic GABA_A_R activation^39^. Instead, what distinguished rested and sleep-deprived cortex was network-independent differences in E_GABAA_, consistent with changes in the contribution of Cl^-^ cotransporter proteins^16,32^. These observations support previous evidence that sleep deprivation can lead to increased contributions of the Cl^-^ importer, NKCC1, and decreased contributions of the Cl^-^ exporter, KCC2^16,32,33^.

The observed changes in E_GABAAR_ are well placed to raise cortical excitability. Periods of sleep deprivation however, are not typically associated with uncontrolled excitation, but more modest changes in network activity^2,7–10,15^. Indeed, following sleep deprivation protocols, animals and humans are typically able to perform tasks that rely upon cortex, albeit at compromised levels^2–6^. Given the adaptive processes available to cortical neurons, it therefore seemed plausible that additional mechanisms could serve to counteract the depolarizing signals in sleep-deprived cortex. In line with this, our *in vivo* current-clamp recordings provide the first evidence that pyramidal neurons engage spike threshold adaptation mechanisms in sleep-deprived cortex, which result in a raised threshold for action potential initiation. Spike threshold adaptation represents a well-established negative feedback mechanism, which operates on a timescale of milliseconds to seconds, is caused by preceding membrane potential effects upon the activation-inactivation and gating kinetics of sodium and potassium channels, and serves to limit excessive excitation^41,42,42–44,50,51,59–61^. Consistent with this, we observed strong relationships between spike threshold and both the preceding absolute membrane potential and rate of membrane depolarization, which are hallmarks of spike threshold adaptation *in vivo*^42^.

Manipulations of spike threshold adaptation are technically challenging. By employing spiking network models, however, we were able to systematically and independently control both the spike threshold and E_GABAAR_ selectively in the simulated pyramidal neurons. This revealed that imposing the E_GABAAR_ and spike threshold values recorded in sleep-deprived cortex was sufficient to generate a net increase in pyramidal neuron spiking and enhance low-frequency oscillatory activity, thereby reproducing experimentally observed *in vivo* effects^16,15,62,10,2,63^. Confirmation of the opposing actions of E_GABAAR_ and spike threshold adaption was demonstrated by varying each of these parameters separately. Altering only E_GABAAR_ resulted in hyperexcitable networks that exhibited evidence of runaway excitation. Meanwhile, altering spike threshold adaptation without the depolarizing shift in E_GABAAR_ resulted in networks that were relatively silent and showed reduced power across the lower frequencies.

In terms of why low-frequency oscillations dominate the SD state, aspects of both a depolarized E_GABAAR_ and spike threshold adaptation can promote synchronous activity at lower frequencies. Depolarizing GABA_A_R signals prolong membrane time constants to promote synchrony on longer timescales^47^, and are less likely to generate higher-frequency oscillatory activity that requires temporally precise hyperpolarizing inhibition^64^. Meanwhile, spike threshold adaption favours frequencies where inter-spike intervals align with the timescale of the adaptation mechanism^65–68^. Stronger and more sustained adaptation tends to synchronize excitatory neurons at lower frequencies^67^. Indeed, multiple factors are likely to contribute to the spike threshold adaptation in sleep-deprived cortex. For example, increases in glutamatergic synaptic activity that follow extended wakefulness would also favour^69–72^. Although aspects of depolarizing GABA_A_R signalling are particularly well placed to promote the more sustained depolarized membrane potentials and slower rates of depolarization that engages spike threshold adaptation. Compared to glutamatergic receptor signalling, phasic GABA_A_R conductances are longer-lasting and extracellular GABA can generate tonic GABA_A_R conductances, where a depolarized E_GABAAR_ would elevate the baseline membrane potential and produce slower changes in depolarization^39,40,73^.

We performed recordings in layer 2/3 pyramidal neurons following a 3-hour period of sleep deprivation at the start of the animal’s inactive period (ZT0-ZT3). It seems likely that the effects upon E_GABAAR_ and spike threshold adaptation, and therefore the interactions between these phenomena, will vary across different neuron types and sleep deprivation protocols. It will therefore be interesting to explore the degree to which these parameters and their interaction can account for reported differences in the effects of extended wakefulness upon different neuron types^20–22,74^. Meanwhile, longer sleep deprivation protocols have been reported to cause sustained changes to intrinsic excitability *in vitro*^75^, which could affect the capacity to observe the more dynamic spike threshold adaptation processes that typically operate on a timescale of milliseconds and seconds.

At a more general level, our findings reveal that periods of sleep deprivation can engage synaptic and intrinsic neuronal mechanisms that oppose one another and thus serve to help maintain overall network stability. In this regard, sleep deprivation could be considered as “using up” adaptive intrinsic mechanisms that would normally be available to cortical neurons. This may mean that when the system is challenged or perturbed in other ways, there is less capacity to regulate excitability and, consequently, the system may transition into a pathophysiological state. Consistent with this idea, animal and human studies have consistently demonstrated that acute sleep deprivation lowers seizure threshold, especially where there is pre-existing seizure predisposition^76–78^. The fact that altered E_GABAAR_ is implicated in a series of neurological disorders that are also associated with sleep disturbances^79,80^, suggests further insights will be gained by considering the interplay between GABA_A_R signalling and adaptive intrinsic properties of cortical neurons.

## Methods

### Animals

All mice were bred, housed and used in accordance with the United Kingdom Animals (Scientific Procedures) Act (1986). For recordings in head-fixed animals, homozygous Gad2-IRES-Cre mice were crossed with homozygous Ai32(RCL-ChR2(H134R)/EYFP) mice (purchased from Jackson Laboratory, Maine, USA) to produce a heterozygous colony expressing channelrhodopsin-2 (ChR2(H134R)-YFP) in Gad2-positive neurons, which includes the main subclasses of GABAergic interneurons. Recordings in freely-moving animals were performed in C57BL/6 wild-type mice (purchased from Charles River, UK). All animals were maintained under the same 12-h:12-h light-dark (LD) cycle for at least 4 weeks to ensure entrainment, were fed ad libitum, and ambient room temperature was maintained at 22 ± 2 degrees Celsius and humidity at 50 ± 20%. Our EEG/EMG recordings and automated vigilance state classification confirmed that animals under these conditions are well entrained, spending an average of 69% of their time asleep during the light (inactive) period and an average of 71% of their time awake during the dark (active) period^16,81^. All experiments were performed in male mice. Female mice were not used in order to minimize potential variance due to the effects of the oestrous cycle upon sleep and circadian processes^82–84^, although we have no reason to expect sex differences in the relationship between spike threshold adaptation mechanisms, depolarizing GABA, and SD. Animals from the same litter were randomly assigned to experimental groups and animal numbers are provided in the figure legends for each experiment.

### Preparation for recordings in head-fixed animals

Head plate fixation was performed under stereotactic surgery on male mice aged 6 weeks. Mice were anaesthetized with isoflurane (Zoetis) and mounted into a stereotaxic frame (Kopf). Subcutaneous analgesia (meloxicam 5mg/kg and buprenorphine 0.1mg/kg) was administered along with intradermal local analgesic (marcain 2 mg/kg) into the scalp. The scalp was shaved (Wahl) and cleaned (Hibiscrub), and eye-protecting ointment (Viscotears) was applied. The scalp was then removed and the site was washed with sterile cortex buffer (containing, in mM: 125 NaCl, 5 KCl, 10 HEPES, 2 MgSO_4_·7H_2_O, 2 CaCl_2_·2H_2_O, 10 Glucose). After drying the skull with adsorbent swabs (Haag-Streit), the periosteum was removed using a micro curette (Fine Science Tools). Tissue adhesive (Vetbond) was applied to secure cranial sutures and to fix the surrounding scalp to the underlying bone. A custom-designed aluminum head plate with a 7 mm well was bonded to the skull, first with adhesive glue (Loctite), and then with serial layers of dental cement (Super-Bond). The well was then covered with silicone sealant (Kwik-Cast). The animal was singly housed and allowed to recover. From day three following head plate fixation, the animal was habituated to head-fixation for increasing time intervals up to 60 minutes. Recordings were performed at 8 weeks of age.

On the day of the recording, animals were studied at Zeitgeber Time 3 (ZT3) having either been allowed to rest for the preceding 3-hours (i.e. between ZT0 and ZT3) as part of their regular sleep pattern (rested, R), or having experienced a 3-hour SD protocol for the preceding 3-hours (sleep-deprived, SD). Head-fixed recordings from animals in the rested and SD conditions were therefore performed at the same time point in the 24-hour cycle. Such an experimental design controls for both the light–dark cycle and circadian processes, enabling us to compare animals at the same point in the circadian cycle but with different sleep–wake histories. The SD protocol involved presenting novel objects to the animal under continuous observation by an experimenter. Once an animal had stopped exploring an object, a new object was presented. This established protocol results in the animal being awake for >98 % of the SD period^16,32^ and, as the SD protocol is performed at the end of the animal’s active period, has been shown to result in increased sleep pressure that is evident in both the awake state and during subsequent sleep^16^. To prepare the brain for recordings, the animal was briefly anaesthetized (10-20 min) with isoflurane, a 0.5 mm craniotomy was created using a dental drill (Foredom), a durectomy performed, and the animal mounted in the head-fixation setup. Recording sessions lasted up to a maximum of 3 hours between ZT3 and ZT6. During the recording session, the experimenter used visual monitoring of the animal and an absence of continuous slow-wave activity in the electrophysiological recordings to confirm that the mouse was awake.

### Gramicidin perforated patch clamp recordings

Gramicidin perforated patch clamp recordings were performed from L2/3 pyramidal neurons in primary visual cortex (V1). The internal pipette solution was prepared immediately prior to recording by combining a high chloride (150 mM) solution (in mM: 141 KCl, 9 NaCl, 10 HEPES) heated to 37 degrees Celsius, with a stock solution of gramicidin A (4 mg/ml - dissolved in dimethyl sulfoxide, DMSO, Merck) to achieve a final concentration of 80 μg/mL gramicidin^85^. The solution was then vortexed (40 s) and sonicated (20 s). Patch pipettes (5-8 MΩ) were back-filled with the gramicidin solution and mounted on a Optopatcher pipette holder (A-M Systems), which contained a 50 μm optic fiber (Thorlabs) connected to a 473nm laser (MBL-FN-473-150mW, CNI Laser). A ground electrode (Multichannel Systems) was placed in the recording well and the patch pipette with positive pressure (200-300 mBar) was inserted into the brain (200-300 μm from the pial surface), the positive pressure lowered to 20-30 mBar, and blind patching commenced.

Once the gigaseal had formed, perforation was then monitored by observing changes in series resistance. Recording protocols were started once the series resistance had stabilized at <100 MΩ. In our experience, recordings that failed to reach these series resistance values within the first 10 minutes never achieved a perforation quality that was suitable for recordings. Rupture or breakthrough of the perforation into whole-cell configuration was detected by a sudden and persistent depolarization of the equilibrium potential of the GABA_A_R (E_GABAAR_), consistent with dialysis of the neuron with the high chloride pipette solution. In a subset of recordings, 15 mM Alexa Fluor 594 (Thermo Fisher) was added to the internal solution so that perforations could also be confirmed by a lack of dye diffusion into the cell, using a two-photon microscope (modified Olympus FV300 confocal scan unit coupled to a Newport Spectra-Physics Mai Tai HP laser, wavelength 820 nm, power 50 mW). In experiments where local network activity was silenced, the AMPA receptor blocker 2,3-dihydroxy-6-nitro-7-sulfamoylbenzo (F) quinoxaline (NBQX) was injected directly into the cortex^86^. The injection pipette contained 100 μM NBQX (Tocris) in ACSF, which was delivered at a rate of 33 nL/min to a total volume of 250 nL.

Whole-cell patch clamp recordings were also performed to enable neurons to be filled with biocytin (internal solution plus 4 mg/ml biocytin), whilst using the same recording criteria as for the gramicidin recordings. To visualize the biocytin-filled neurons, brains were fixed via transcardial perfusion of phosphate buffered solution (PBS, 0.1 M) and 4 % paraformaldehyde solution (PFA), stored for 24 h at 4°C in 4 % PFA, and then washed and stored in PBS containing 0.05 % sodium-azide. Brains were sectioned into 100 μm thick coronal slices in PBS (HM650V microtome, ThermoScientific) and sections were incubated in Streptavidin-Cy3 (1:1000, Thermo Fisher) for 2 h at RT, before being mounted with Vectashield (Vectorlabs) onto glass slides (Avantor) and imaged using a LSM 880 microscope (Zeiss). All images were captured using a 20x water-immersion objective (W Plan-Apochromat NA 1.0) using ZEN software (Zeiss) and images were processed in ImageJ software (NIH).

### Data acquisition

Patch clamp recordings were performed using a Multiclamp 700B amplifier (Molecular Devices) and digitized at 20 kHz (Digidata 1550, Molecular Devices). A HumBug noise eliminator (Digitimer) was used to remove 50 Hz noise. Membrane and recording properties were calculated by measuring the change in current in response to a -10 mV step during voltage-clamp recordings. The series resistance (R_s_) was calculated from both the peak current elicited by the -10 mV voltage step, and by estimating the peak after fitting an exponential to the decay of the current transient response to the -10 mV step. These methods gave similar values and so the numerical average was used as a final estimate of R_s_. The mean R_s_ values for the four conditions were: rested, 55 ± 2.8 MΩ; SD, 58 ± 1.3 MΩ; rested+NBQX, 50 ± 2.5 MΩ; SD+NBQX, 56 ± 3.3 MΩ. To calculate the membrane resistance (R_m_), R_s_ was subtracted from the measured input resistance. Online R_s_ compensation was not used, as the high amounts of activity in the *in vivo* brain would cause large fluctuations in input current, which increases the rate of perforation rupture^87,88^. Therefore, to correct for R_s_ effects, offline correction was performed.^89^ The voltage drop caused by the series resistance (R_s_) was calculated by multiplying the measured current response with the R_s_. The voltage drop was then subtracted from the command voltage to estimate the neuron’s membrane potential.

### Measuring synaptic E_GABAAR_ and DF_GABAAR_

A voltage ramp protocol was used to measure synaptic E_GABAAR_. This involved clamping the neuron at -70 mV, and then imposing two consecutive voltage ramps, each lasting 150 ms, which extended from 60 mV below the holding voltage to 40 mV above the holding voltage (i.e. from -130 mV to -30 mV, at a rate of 0.7 mV/ms). The first ramp (i.e. the ‘baseline’ ramp) sampled the neuron’s intrinsic membrane currents and the second ramp (i.e. the ‘light’ ramp) included a light-evoked synaptic GABA_A_R conductance. The light-evoked synaptic GABA_A_R conductance was elicited with a 10 ms light pulse that coincided with the start of the ramp, to ensure the evoked GABA_A_R current was at its peak during the ramp^39,90^. The currents from both ramps were then superimposed on a current-voltage (IV) plot using the series corrected membrane potentials and, to avoid action potentials and capacitance transients, the current responses were cropped to only include regions that were clear of these sources of contamination. A straight line was fitted to both currents. The current from the baseline ramp was used to infer the equilibrium potential of the holding current (from which the resting membrane potential, RMP, could be inferred), defined as the voltage at which the fitted line was equal to zero. The point at which the fitted lines for the two ramps intersected was defined as synaptic E_GABAAR_. The synaptic DF_GABAAR_ was calculated by subtracting the measured synaptic E_GABAAR_ from the RMP.

### Membrane potential dynamics and spike threshold adaptation

Current-clamp recordings were used to quantify membrane potential dynamics and spiking properties. To quantify subthreshold V_m_ dynamics, probability density function plots were created by combining all membrane potential values (after excluding action potentials) for all neurons in a particular experimental condition, and fitting the resulting data with a Gaussian kernel-density. To infer the state of local network activity, the level of subthreshold synaptic activity was calculated by measuring the rate of change in the V_m_ over time (after excluding action potentials). The rate of change in V_m_ was calculated by winsorizing the first derivative (V_m_ dV/dt in mV/ms), and calculating the mean of the modulus of the V_m_ dV/dt for each neuron. To quantify action potential properties, including spike threshold adaptation, phase plots were generated to show the rate of change in V_m_ (dV/dt) as a function of V_m_. Spike threshold (Spike_Thres_) was defined as the V_m_ at the onset of an action potential, where onset was defined as the first point at which dV/dt crossed one standard deviation above the mean dV/dt calculated over a 2 ms window that encompassed the spike. The mean V_m_ and rate of V_m_ depolarization before each spike were measured over a 2 ms window immediately preceding the spike.

### LFP recordings in freely-moving mice

For *in vivo* LFP recordings in freely-moving animals, vigilance states were determined by monitoring the electroencephalogram (EEG) and electromyogram (EMG). Implantation of the recording devices was performed under stereotactic surgery, aseptic technique, isoflurane anaesthesia (3-5% for induction and 1-2% for maintenance) and constant body temperature monitoring. Analgesia (buprenorphine and meloxicam) was provided at the beginning of surgery and during recovery. Craniotomies were performed over left somatosensory cortex (AP -1.2 mm, ML -3 mm from Bregma) for the LFP electrode (insulated untwisted tungsten wire, 0.125 mm; C315G-MS303/2, PlasticsOne), over the right frontal cortex (AP +2 mm, ML +2 mm) and right occipital cortex (AP −3.5 mm, ML +2.5 mm) for EEG electrodes, and over the cerebellum (-1.5 mm posterior from Lambda, ML 0) for the reference electrode (all stainless steel screw electrodes, Fine Science Tools). For the EMG, two wire electrodes were inserted into the left and right neck muscles, with one signal acting as reference to the other. The LFP electrode, EEG and reference screws, plus EMG electrodes were connected to an eight-pin surface mount connector (8415-SM, Pinnacle Technology). Implants were secured with a non-transparent dental cement (Super-Bond), and animals were allowed to recover for at least 1 week before recordings. Animals were moved to a recording chamber and housed individually in a Plexiglas cage (20.3 x 32 x 35 cm). Recordings were performed using a 128-channel Neurophysiology Recording System (Tucker-Davis), acquired using the electrophysiological recording software, Synapse (Tucker-Davis), and stored locally for offline analysis. Signals were continuously recorded and stored at a sampling rate of 305 Hz, then resampled at 256 Hz and converted into the European Data Format. Data were processed for automated vigilance scoring using the open-source sleep stage classifier, Somnotate^16,81^. Rested and SD LFPs were compared on consecutive days in the same mouse. The rested LFP was collected on the first day, whilst the animal was allowed to follow its normal sleep-wake pattern, and throughout the light period (ZT0-ZT12) to ensure sufficient sampling of wake periods. On the second day, the mouse was subjected to the same SD protocol as for the head-fixed recordings. This involved a 3-hour SD protocol at the start of the animal’s inactive period (ZT0-ZT3), and the SD LFP was collected during the third hour of the SD protocol (ZT2-ZT3). No distinctions were made within the waking state (e.g. quiet versus active waking)^17^.

### Neuronal network simulations

Computational models were used to explore the effects of spike threshold adaptation, E_GABAAR_, and RMP on glutamatergic neuron spiking activity and the LFP. The power spectrum of excitatory synaptic inputs was used as a proxy of the LFP^53^. Networks were constructed using the neuron simulator Brian 2^91^ and comprised 1000 glutamatergic pyramidal neurons and 250 GABAergic interneurons. Each neuron was modelled as a single compartment, current-based leaky integrate-and-fire neuron. Constant model parameters were set as shown in **Table 1**.

**Table 1:** Constant model parameters for neuronal network modelling.

|  |  |
| --- | --- |
| Membrane capacitance | 200 pF |
| Leak conductance (nS) | 10 nS |
| Glutamatergic synapse reversal potential | 0 mV |
| Glutamatergic synapse time constant | 5 ms |
| Glutamatergic neuron refractory period | 5 ms |
| GABAergic synapse time constant | 10 ms |
| GABAergic neuron refractory period | 1 ms |

The glutamatergic neurons connected to each other with a probability of 10 %^92^; Glutamatergic to GABAergic neurons, GABAergic to glutamatergic neurons, and GABAergic neurons with each other, all connected with a probability of 60 %^93^. All synaptic weights were initialized to 1 nS, but at the start of each simulation, homeostatic inhibitory synaptic plasticity was used to establish a balance of excitatory and inhibitory inputs to each neuron^94^, according to defined plasticity parameters (STDP time constant of 20 ms, postsynaptic neuron target firing rate of 20 Hz), and then synaptic weights were frozen. To create four different network conditions, the spike threshold, E_GABAAR_, and RMP for glutamatergic neurons was varied according to our experimentally-observed values, as shown in **Table 2**. The corresponding parameters for GABAergic neurons remained constant at -50 mV, -60 mV and -60 mV, respectively.

**Table 2:** Glutamatergic neuron parameters used to create four network conditions.

| | Condition 1<br>“R”<br>(rested) | Condition 2<br>“SD”<br>(sleep-deprived) | Condition 3<br>“ST-”<br>(SD w/o spike<br>thres. adapt.) | Condition 4<br>“EG-”<br>(SD w/o $E_{GABAAR}$<br>plasticity) |
| --- | --- | --- | --- | --- |
| Spike threshold | -52.3 mV | -45.5 mV | -52.3 mV | -45.5 mV |
| $E_{GABAAR}$ | -62.2 mV | -52.1 mV | -52.1 mV | -62.2 mV |
| RMP | -63.2 mV | -58.7 mV | -58.7 mV | -58.7 mV |

Each neuron received a noise input current and a shared input current. The noise input was modelled as random independent values drawn from log-normal distributions with the scale parameter mu set to zero, and the shape parameter sigma set to 1:

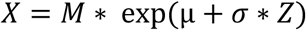

with Z being a standard normal variable. Meanwhile, the shared input current followed a 1/f power spectrum, as widely observed in LFP and EEG recordings^95^. This current was generated using the ‘powerlaw_psd_gaussian’ function from the colorednoise python module. The exponent beta was set to one, resulting in a pink noise distribution of shared input current values. All other parameters were kept at default values. Both the noise and shared input currents were shifted and rescaled to have an expected mean and standard deviation of 50 pA, and were varied every 10 ms. Networks were first balanced for 30 s in the SD condition, and then simulated for a period of 300 s. Each network simulation was repeated 20 times.

### Statistical analysis

All data is reported as mean ± standard error of the mean (SEM). For comparative statistics, the distribution of the continuous data was first established using the Shapiro-Wilk test for normality, which guided the subsequent use of appropriate parametric and non-parametric tests. All statistical analyses were performed using the Python SciPy library. All tests were two-tailed, with a confidence level of at least a 95%. Statistical tests were unpaired unless stated. Details of the data values, sample sizes and statistical measurements are provided in the figure legends.

## Supplementary figures

**Supplementary Figure 1:**
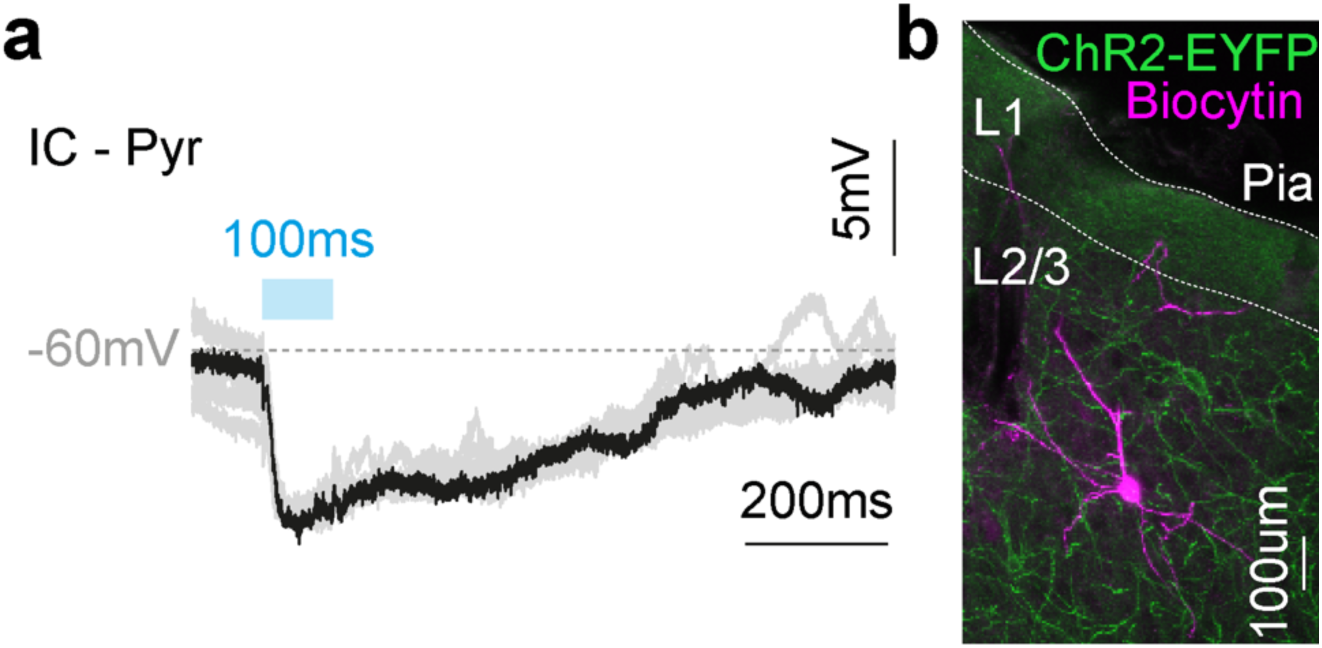
Confirming targeting of L2/3 pyramidal neurons. **a**, Example light-evoked postsynaptic responses (100 ms, blue light pulse) recorded in current-clamp mode from a neuron in the gramicidin perforated patch clamp configuration *in vivo*. As ChR2 was expressed only in cortical interneurons, the lack of a direct light-evoked spike or membrane depolarization, the presence of a light-evoked fast postsynaptic inhibitory response, and a recording depth relative to the pial surface that corresponded to L2/3, were the criteria used to identify L2/3 pyramidal neurons for further analysis. **b**, Confocal image of a biocytin-filled L2/3 pyramidal neuron stained with streptavidin-fluorophore conjugate, streptavidin-CY3. A series of whole-cell patch clamp recordings were performed to enable the recorded neuron to be filled with biocytin, using the same criteria as the gramicidin recordings. The morphology of the neurons was imaged post-mortem and provided further confirmation of targeting L2/3 pyramidal neurons.

**Supplementary Figure 2:**
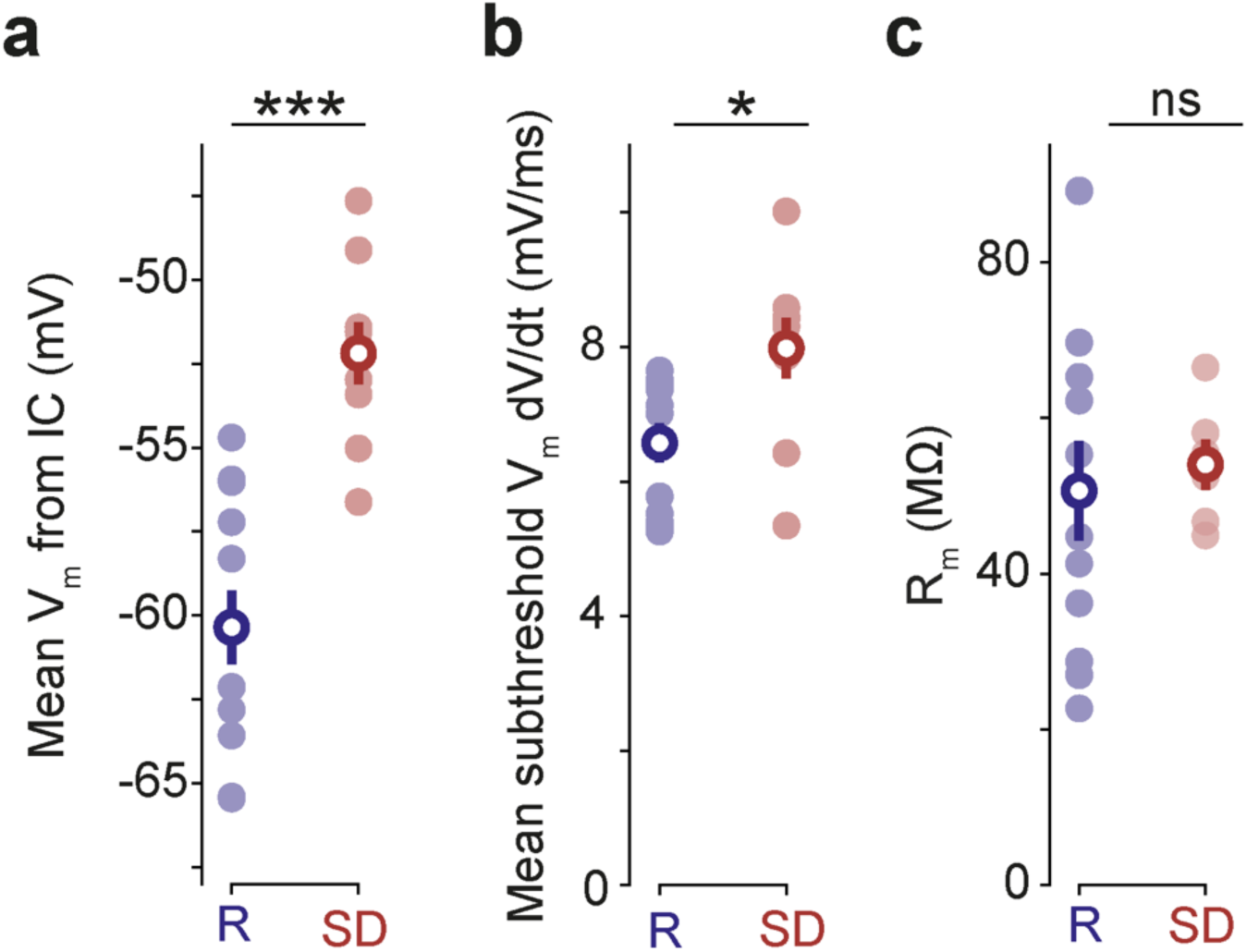
L2/3 pyramidal neurons report increased levels of network activity in the sleep-deprived cortex. **a**, Current-clamp recordings revealed that L2/3 pyramidal neurons exhibit more depolarized mean membrane potentials (V_m_) in sleep-deprived (SD, red) mice compared to rested (R, blue) mice (R: -60.3 ± 1.1 mV, n = 12 neurons from 10 mice, vs. SD: -52.2 ± 0.9 mV, n = 9 neurons from 9 mice, p < 0.001, *t-test*). **b**, Neurons in SD mice exhibited larger fluctuations in subthreshold V_m_ (R: 6.6 ± 0.3 mV/ms, n = 12 neurons from 10 mice, vs. SD: 7.9 ± 0.5 mV/ms, n = 9 neurons from 9 mice, *p < 0.02, t-test*). **c**, Membrane resistance (R_m_) was comparable between neurons in rested and SD mice (R: 50.6 ± 6.4 MΩ, n = 13 neurons from 10 mice vs. SD: 54.1 ± 3.2 MΩ, n = 6 neurons from 6 mice, p = 0.74, *t-test*). ‘ns’, non-significant; p > 0.05; *, p < 0.05; ***, p < 001.

**Supplementary Figure 3:**
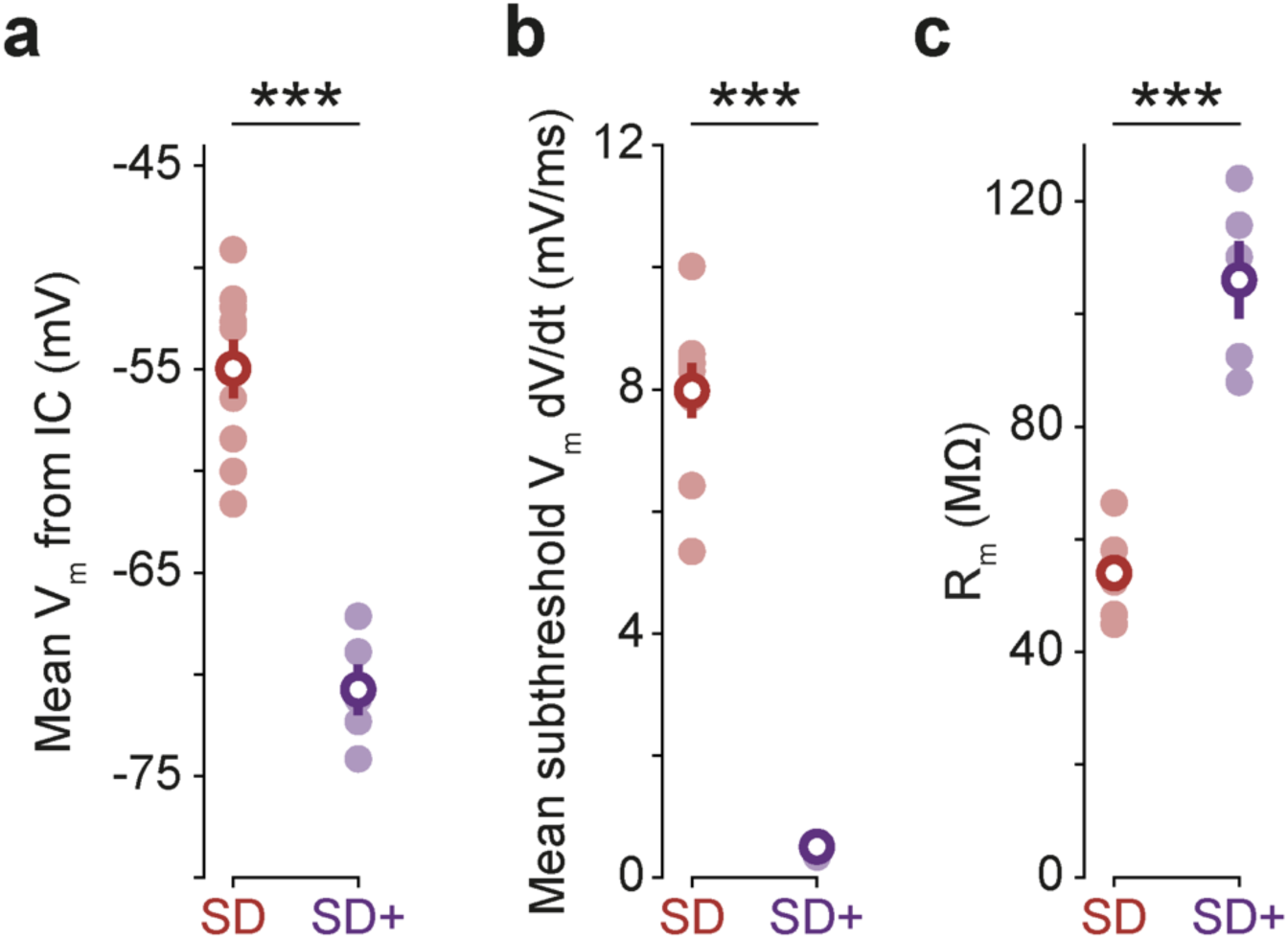
Blocking glutamatergic signaling silences local network activity in the sleep-deprived cortex. **a**, Current-clamp recordings revealed that neurons in sleep-deprived (SD, red) mice exhibit more hyperpolarized mean membrane potentials (V_m_) following the local delivery of NBQX (SD+, purple) (SD: -54.9 ± 1.4 mV, n = 9 neurons from 9 mice, vs. SD+: -70.8 ± 1.2 mV, n = 5 neurons from 5 mice, p < 0.001, *t-test*). **b**, Neurons in SD mice exhibited an almost complete suppression of subthreshold V_m_ fluctuations following the local delivery of NBQX (SD+), consistent with a silencing of local network activity (SD: 7.9 ± 0.5 mV/ms, n = 9 neurons from 9 mice, vs. SD+: 0.5 ± 0.04 mV/ms, n = 5 neurons from 5 mice, p < 0.001, *t-test*). **c**, Neurons in SD mice exhibited an increase in membrane resistance (R_m_) following the local delivery of NBQX (SD+) (SD: 54.1 ± 3.2 MΩ, n = 6 cells, 6 mice vs. SD+: 106.1 ± 6.9 MΩ, n = 5 cells, 5 mice, p < 0.001, *t-test*). ***, p < 0.001.

**Supplementary Figure 4:**
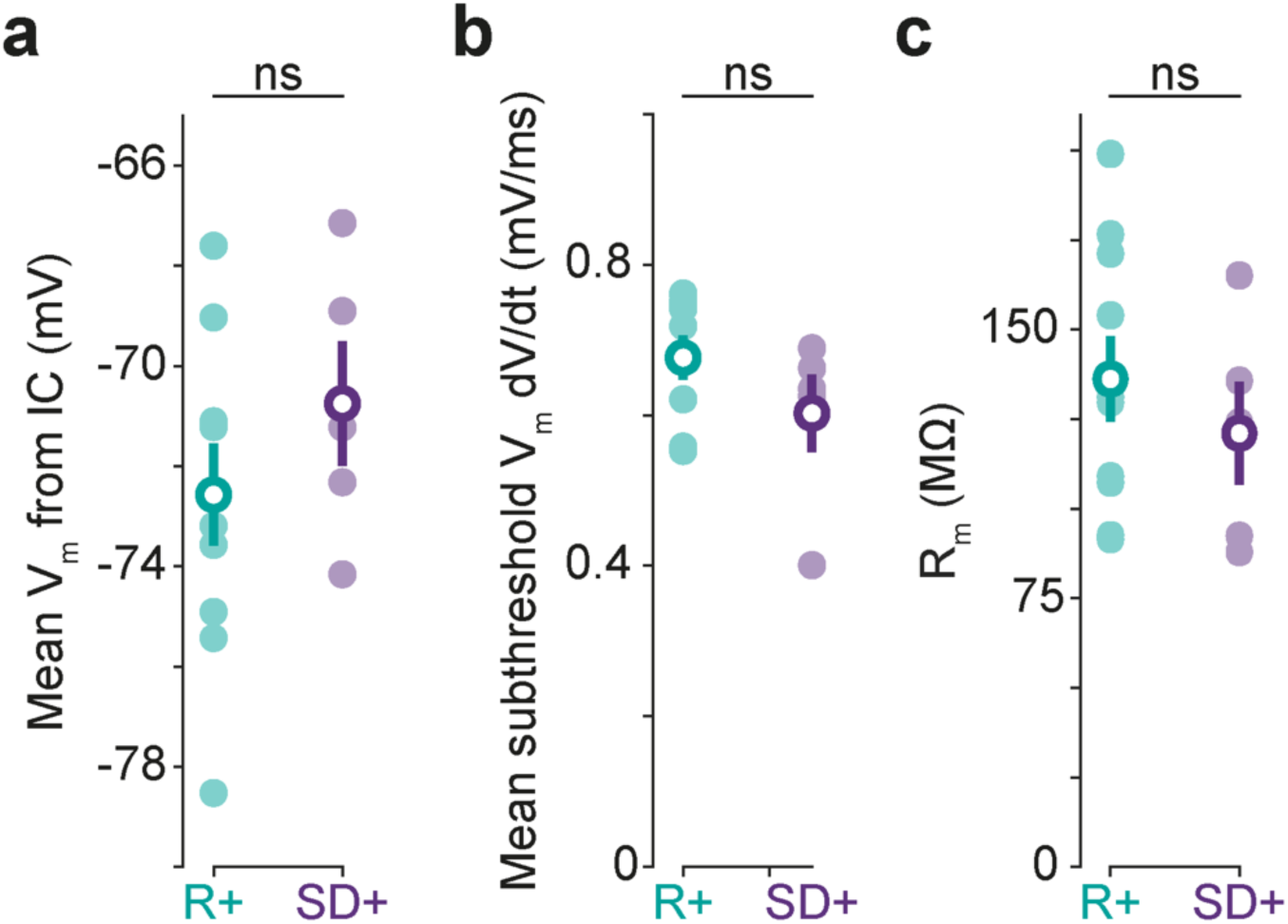
Blocking glutamatergic signaling silences and equalizes local network activity in rested and sleep-deprived cortex. **a**, Current-clamp recordings revealed that the local delivery of NBQX in rested (R+, turquoise) and sleep-deprived (SD+, purple) mice resulted in comparable mean membrane potentials (V_m_) (R+: -72.6 ± 1.0 mV, n = 10 neurons, 7 mice, vs. SD+: -70.8 ± 1.2 mV, n = 5 neurons, 5 mice, p = 0.3, *t-test*). **b**, The local delivery of NBQX resulted in an almost complete suppression of subthreshold V_m_ fluctuations in rested and SD mice, consistent with a silencing and equalizing of local network activity (R+: 0.68 ± 0.03 mV/ms, n = 10 neurons, 7 mice, vs. SD+: 0.60 ± 0.05 mV/ms, n = 5 neurons, 5 mice, p = 0.2, *t-test*). **c**, The local delivery of NBQX resulted in comparable membrane resistance (R_m_) in rested (R+, turquoise) and SD (SD+, purple) mice (R+: 136.2 ± 11.9 MΩ n = 10 neurons, 7 mice vs. SD+: 121.1 ± 14.3 MΩ, n = 5 neurons, 5 mice, p = 0.5, *t-test*). ‘ns’, non-significant.

**Supplementary Figure 5:**
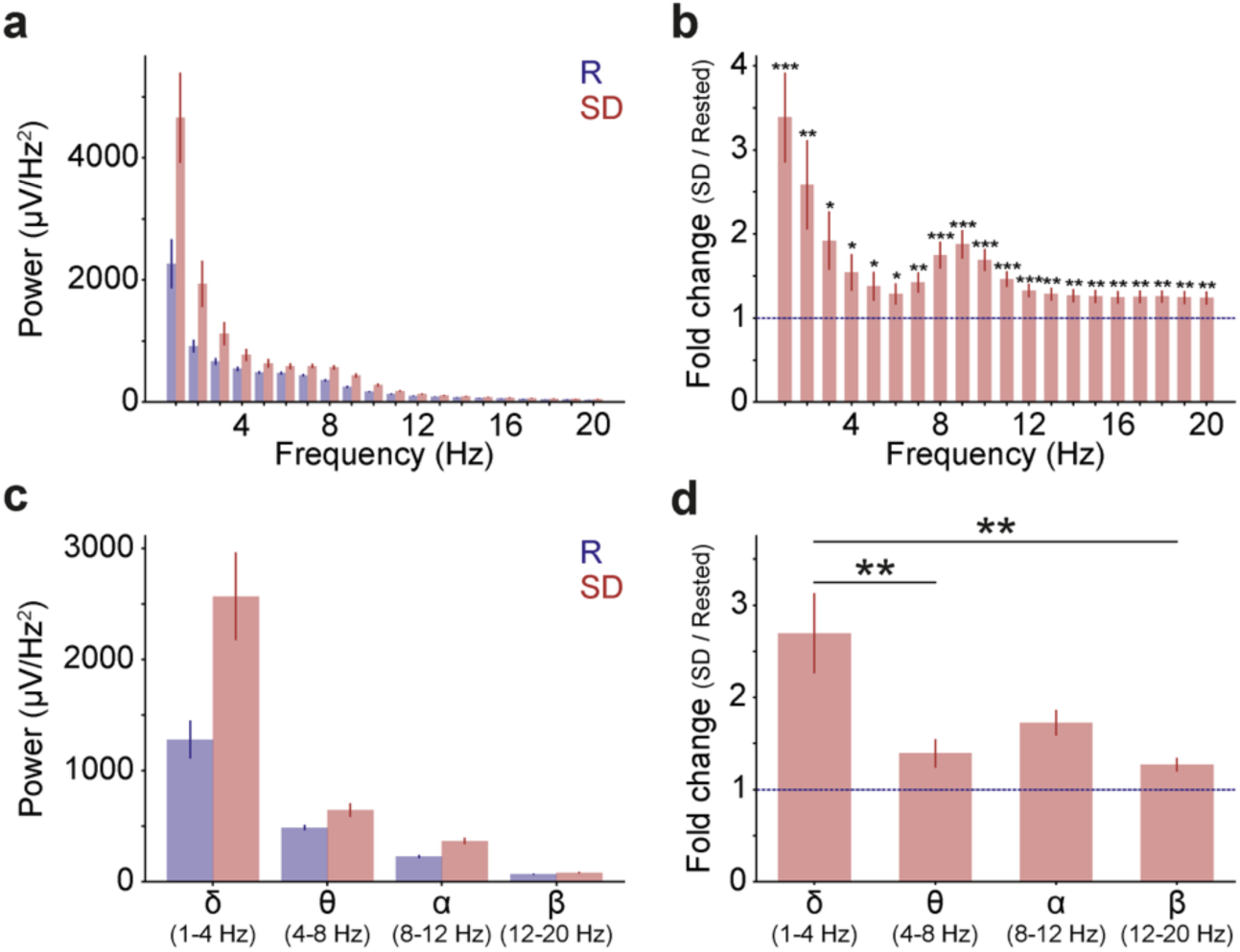
Experimentally-observed changes in local field potentials recorded from sleep-deprived cortex. **a**, Local field potential (LFP) power per 1-Hz frequency bin for mice in their rested (R, blue) and sleep-deprived (SD, red) state (n = 22 mice, p < 0.001, *two-way repeated measures Anova*). **b**, Fold-change in LFP power per 1-Hz frequency bin (SD/Rested). Values above 1 indicate an increase in the SD state, asterisks indicate statistical differences from a value of 1 (dashed blue line; *one-sample t-tests*). **c**, LFP frequency band-specific power comparisons for mice in their rested (R, blue) and sleep-deprived (SD, red) state. **d**, Fold-change in LFP band power (SD/Rested) showed a main effect of frequency band (p < 0.001, ANOVA), asterisks indicate differences relative to the delta frequency band (*Bonferroni post-hoc tests*). *, p < 0.05; **, p < 0.01; ***, p < 0.001.

**Supplementary Figure 6:**
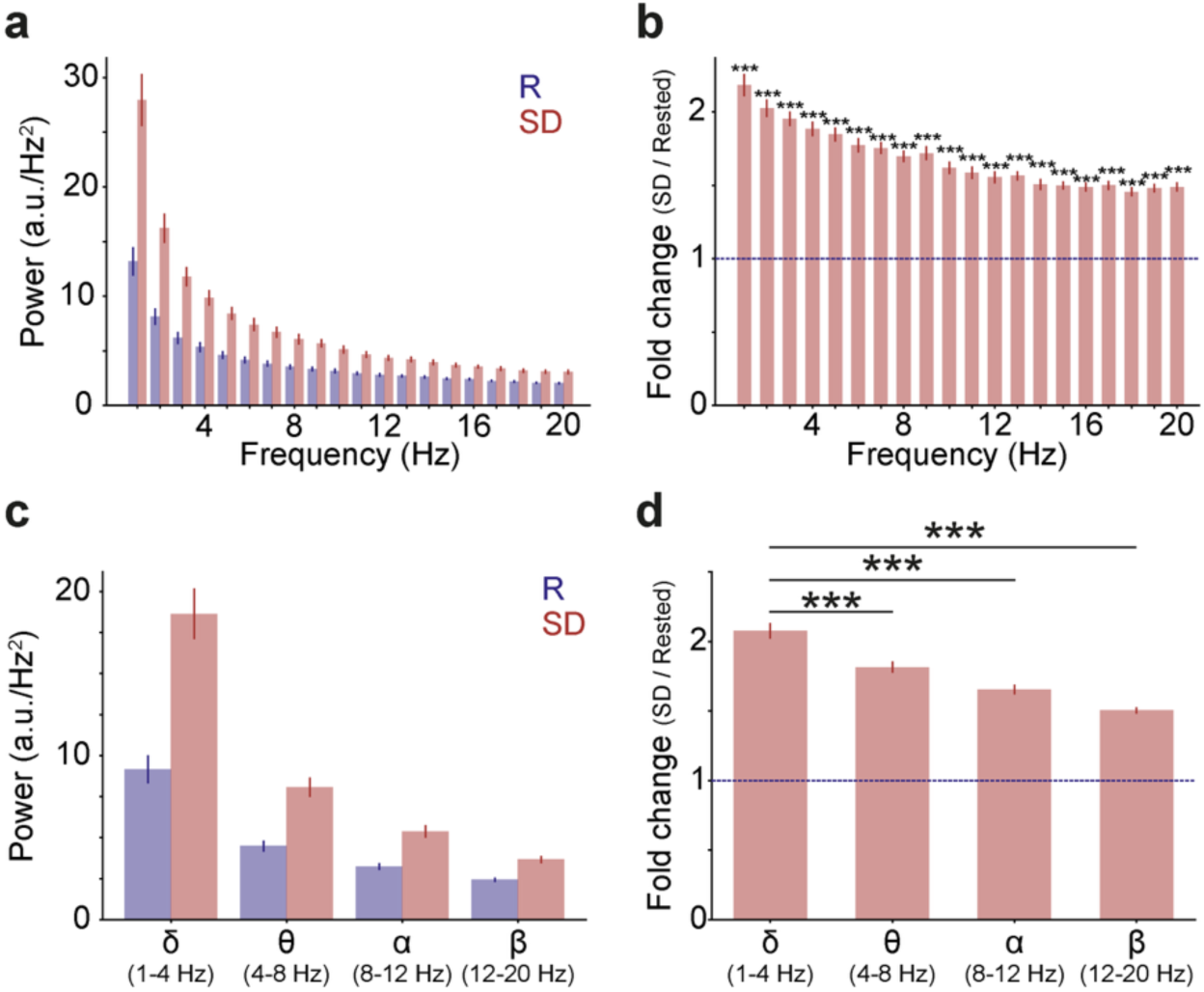
Changes in local field potentials in a simulated sleep-deprived network. **a**, Model-derived local field potential (LFP) power for each 1-Hz frequency bin in rested (blue) and sleep-deprived (SD; red) simulated networks (n = 20 simulations per condition, p < 0.001, *two-way repeated measures Anova*). **b**, Fold-change in LFP power per 1-Hz frequency bin (SD/Rested). Values above 1 indicate an increase in the SD network, asterisks indicate statistical differences from a value of 1 (dashed blue line; one-sample t-tests). **c**, LFP frequency band-specific power comparisons for rested (R, blue) and sleep-deprived (SD, red) networks. **d**, Fold-change in LFP band power (SD/Rested) showed a main effect of frequency band (p < 0.001, ANOVA), asterisks indicate differences relative to the delta frequency band (Bonferroni post-hoc tests). *, p < 0.05; **, p < 0.01; ***, p < 0.001.

## Acknowledgments

We would like to thank members of the Akerman lab for advice and comments. We thank Lex Kravitz, Luigi Petrucco and Ethan Tyler for sharing their mouse illustrations on the Sci-Draw opensource platform. The research leading to these results has received funding from the European Research Council under grant agreement 617670, MRC project MR/Z504567/1, MRC project MR/S01134X/1, BBSRC project BB/S007938/1, and a Research Grant from the Leverhulme Trust. In addition, this work was supported by a Shaun Johnson Memorial Scholarship sponsored by the Leverhulme Trust and Mandela Rhodes Foundation (RJB), and a Sir Henry Wellcome Postdoctoral Fellowship 206500/Z/17/Z (HA).

## Author contributions

RB, HA and CJA conceptualized the project. RB, HA, VVV and CJA designed the experiments. RJB performed and analysed the *in vivo* patch clamp recordings. HA performed the LFP recordings. PJNB designed and performed the neuronal network modelling. RB and CJA wrote the manuscript with input from all authors. CJA supervised the project.

## Data availability

Source data for all Figures is included. Raw data from electrophysiological recordings are available from the corresponding authors upon reasonable request.

## Code availability

Custom-made code used for analyses is available from the corresponding authors upon reasonable request. The code used to run neuronal network simulation can be accessed at: https://gist.github.com/paulbrodersen/ea306bee72c4ea7e9eada8cb90e79e78

**Competing interest:**

The authors declare no competing interests.

